# Can the Kuznetsov Model Replicate and Predict Cancer Growth in Humans?

**DOI:** 10.1101/2022.02.02.478884

**Authors:** Mohammad El Wajeh, Falco Jung, Dominik Bongartz, Chrysoula Dimitra Kappatou, Narmin Ghaffari Laleh, Alexander Mitsos, Jakob Nikolas Kather

**Author notes:** Shared last authorship and corresponding authors,.

## Abstract

Several mathematical models to predict tumor growth over time have been developed in the last decades. A central aspect of such models is the interaction of tumor cells with immune effector cells. The Kuznetsov model (Kuznetsov et al. (1994), Bulletin of Mathematical Biology, vol. 56, no. 2, pp. 295–321) is the most prominent of these models and has been used as a basis for many other related models and theoretical studies. However, none of these models have been validated with large-scale real-world data of human patients treated with cancer immunotherapy. In addition, parameter estimation of these models remains a major bottleneck on the way to model-based and data-driven medical treatment. In this study, we quantitatively fit Kuznetsov’s model to a large dataset of 1472 patients, of which 210 patients have more than six data points, by estimating the model parameters of each patient individually. We also conduct a global practical identifiability analysis for the estimated parameters. We thus demonstrate that several combinations of parameter values could lead to accurate data fitting. This opens the potential for global parameter estimation of the model, in which the values of all parameters are fixed for all patients. Furthermore, by omitting the last two or three data points, we show that the model can be extrapolated and predict future tumor dynamics. This paves the way for a more clinically relevant application of mathematical tumor modeling, in which the treatment strategy could be adjusted in advance according to the model’s future predictions.

## 1. Introduction

Cancer immunotherapy with immune checkpoint inhibitors has revolutionized the treatment of patients with solid tumors in the last ten years. In addition to chemotherapy and molecularly-targeted therapy, immunotherapy provides a new set of tools for the oncology toolkit [1]. In several tumor types such as melanoma, non-small cell lung cancer (NSCLC), and genito-urinary tumors, immunotherapy has markedly improved the average life expectancy of patients with advanced disease. Both laboratory and clinical experiments have verified the importance of the immune system in fighting cancer [2]–[4]. Patients who suffer from acquired immunodeficiency syndrome (AIDS) are very susceptible to having some rare forms of cancer [2], [5]. This also shows the significant role the immune system plays against cancer.

One of the fundamental problems in treating patients with cancer immunotherapy is the lack of predictive biomarkers. Ideally, before the treatment begins, patients could be selected for immunotherapy, but existing biomarkers fail to deliver a high predictive value in most tumor types [6]. In addition, most patients who initially respond to immunotherapy progress experience a relapse: the tumor later on develops immune escape mechanisms due to evolutionary pressure. Forecasting the time of relapse or treatment resistance is of high practical relevance [7], [8]. However, predictions of such changes in the tumor behavior are currently not possible in clinical routine. The main problem is that most biomarkers such as tumor mutational burden (TMB) are static, i.e. they are measured at a given time point but are not dynamically updated as the tumor evolves.

In other complex systems such as financial markets [9], climate systems [10] or complex industrial processes [11], differential equation models can provide a prediction of the behavior of the system over time. By analogy, in oncology, a number of mathematical models to predict tumor growth over time have been developed in the last decades [12]. Most notably, multiple of these models explicitly include the interactions of tumors with the immune system and are therefore in principle suited to model response and resistance to cancer immunotherapy [13]. Boer and Hogeweg (1986) modeled the cellular immune reaction to tumors. They demonstrated that small doses of antigens lead to tumor dormancy [2], [14]. Kirschner and Panetta (1998) linked the oscillations in the tumor size and the long-term tumor regression to the dynamics among immune cells, tumor cells, and Interleukin-2 [2], [15]. The most prominent of these models was presented by Kuznetsov et al. [16] in 1994. Kuznetsov’s model has served as a blueprint for many other related models [2], [17], [18] and has been investigated in several theoretical studies [2], [19]–[21].

However, none of these established oncological models are currently being used in the clinic. What is more, very few of these models have been systematically fitted to actual clinical data. While some studies have fitted models to murine tumors on a small scale [2], [22], [23], the pronounced differences between mice and humans preclude the transfer of such insights to real-world cancer patients [24].

The structures of these mathematical models are well defined [16], [25]–[27]. However, in the complex biological environment of cells, little is known about the associated parameters and kinetic constants. The parameter values are essential for quantitative modeling and prediction of cancer progression. In mechanistic models, one can integrate the data from various experimental procedures and sources, and design in-silico experiments to generate hypotheses for underlying mechanisms [28], [29]. By fitting the model to the experimental data, we reverse-engineer the parameters of the system. Parameter estimation of mathematical cancer models remains a major bottleneck on the way to model-based and data-driven medical treatment of the future.

In this study, we use a mathematical model based on Kuznetsov’s model to characterize the interactions between the growing tumor and the immune system, and aim to fill this conceptual gap in the literature. We use a large dataset of thousands of cancer patients who underwent cancer immunotherapy as part of clinical trials. We then investigate how well the model can represent the actual tumor volume changes over time in these patients. After estimating the parameters of the model, we conduct an identifiability analysis to examine the uniqueness of the estimated parameters (i.e., whether we have over-fitting). Finally, we investigate if the model can be used to forecast treatment response or relapse under immunotherapy.

The remainder of the article is structured as follows. First, we briefly present the acquisition of patients data and its pre-treatment in Section 2. In Section 3, we then provide the model and all the methods used to estimate model parameters and fit the data, conduct parameter identifiability analysis and extrapolate the model for tumor growth prediction. Finally, before drawing conclusions in Section 5, we present and discuss the obtained results in Section 4.

## 2. Data Acquisition

We briefly provide here the declaration and sources of the experimental data. For more details, please refer to [30].

### 2.1. Declarations and Data Sharing

We followed the Declaration of Helsinki and International Ethical Guidelines for Biomedical Research Involving Human Subjects developed by the Council for International Organizations of Medical Sciences (CIOMS). In this study, we used a publicly available set of anonymized patient data shared by [30], which is originally derived from five large clinical trials, as we describe below. Patients gave their informed consent for data analyses as part of the original clinical trials. No specific ethical approval was sought or required for this retrospective analysis of a publicly available dataset.

### 2.2. Data Measurement and Pre-treatment

The original data for this study is from five clinical trials which were designed to evaluate the efficiency of Atezolizumab (an immune checkpoint inhibitor). Table 1 shows the original number of patients and their treatment arms. Four out of these five trials evaluated the effect of Atezolizumab on NSCLC and cohort GO29293 reported this efficiency on bladder cancer. In two of the cohorts (GO28753, GO28915), patients’ responses to Atezolizumab treatment were compared to the outcome of the second treatment arm who received Docetaxel (a chemotherapy drug) as treatment. In all the trials, the longest and shortest diameter (LD and SD) of the target and non-target lesions (measured manually based on the CT scans) alongside the time intervals are reported. In this study, we used the anonymized and publicly available subset of data from [30]. This subset is created by selecting the patients with three or more measurement points. For each of these patients, only the LD measurement for one target lesion has been selected. For this reason, the total number of original patients of clinical trials has been decreased from 2693 to 1472. All the pre-processing details for the data generation has been described in more details in [30]. Moreover, before using the data we pre-processed it by first removing repetitive and null inputs. Following [30]–[32], we converted the measured LD in mm to the number of tumor cells (TC) by considering 4/3 × (8 × 10^−6^) mm^3^/TC. We also considered only patients with more than six net measurements. Figure 1 shows the number of patients per study and arm before and after data pre-processing. Finally, the total number of patients has been reduced from 1472 to 210.

**Table 1:** Description of the original data.

| Study ID | Cancer Type | No. of Patients | Treatment |
| --- | --- | --- | --- |
| GO28625 [33] | NSCLC | 138 | Atezolizumab |
| GO28753 [34] | NSCLC | 287 | Atezolizumab/Docetaxel |
| GO28754 [35] | NSCLC | 657 | Atezolizumab |
| GO28915 [36] | NSCLC | 1182 | Atezolizumab/Docetaxel |
| GO29293 [37] | Bladder Cancer | 429 | Atezolizumab |

**Figure 1:**
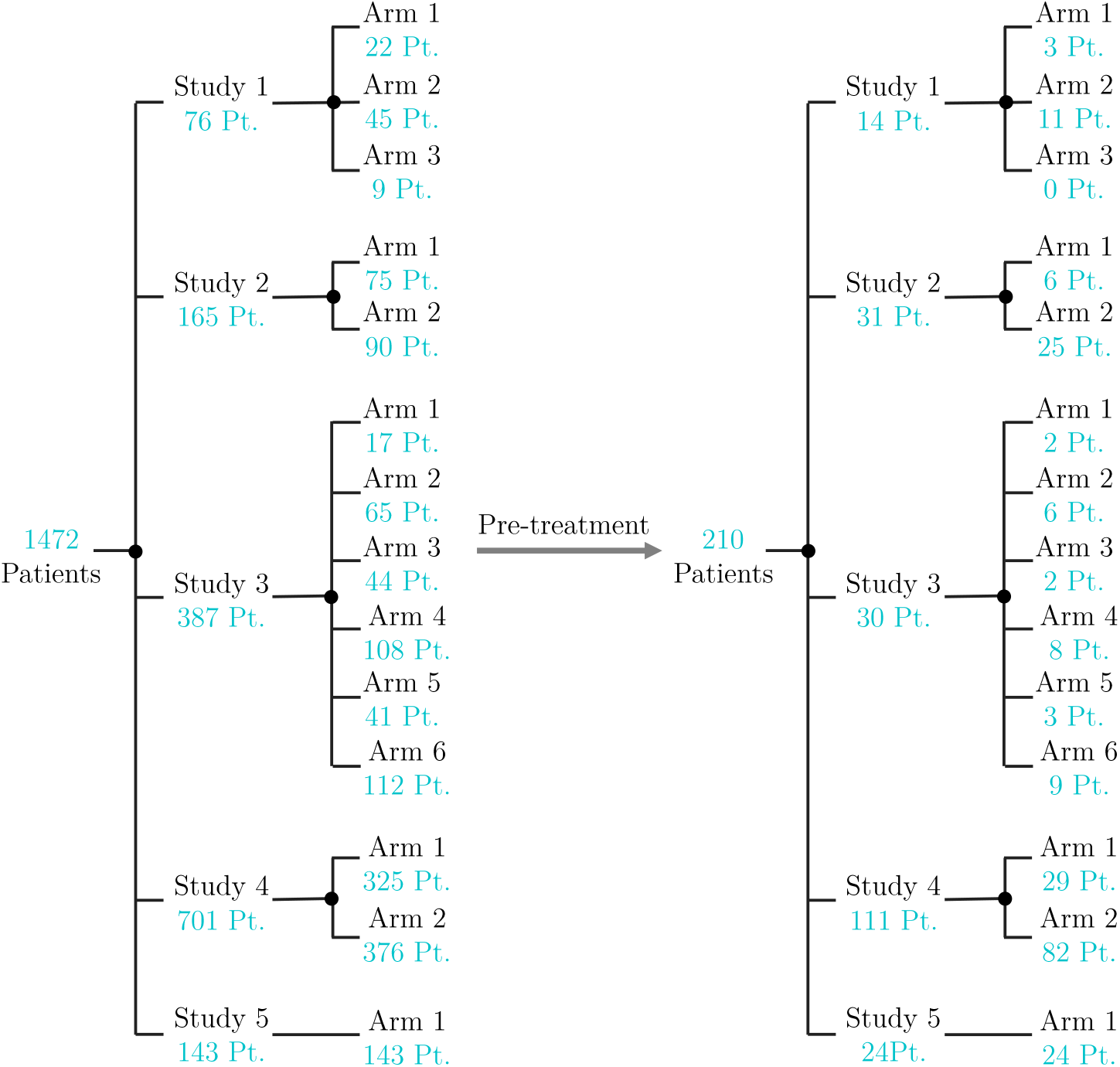
Number of patients (Pt.) considered per studies and arms, before and after data pre-processing.

## 3. Methods

In this section, we introduce the mathematical model of Kuznetsov [16] with slight modifications. Afterwards, we show the optimization formulations for: the estimation of the model parameters by fitting the clinical data and for the parameter identifiability analysis. We finally investigate the extrapolation capabilities of the model.

### 3.1. Mathematical Tumor Model

To predict the frequently observed phenomena in clinics like tumor dormancy and tumor size oscillation, the tumor mathematical model has to include terms related to the response of the immune system. The inclusion of the entire immune system in the mathematical model can be very difficult [38]. The anti-tumor immune response has highly nonlinear dynamics which are complicated and not well understood. Therefore, models that describe the immune system response to tumor presence should necessarily focus on those elements of the immune system that have the highest effects on tumor dynamics [2]. Kuznetsov’s model [16] describes the response of the cytotoxic T lymphocyte (CTL) to the growth of an immunogenic tumor. Usually, a cell-mediated immune response to a tumor takes place. The cytotoxic T lymphocytes and natural killer (NK) cells play the main role. The model considers immunogenic TC that are attacked by cytotoxic effector cells (EC). The EC can be, for example, CTL or NK cells. The model takes into account the possibility of EC inactivation as well as the infiltration of the TC by EC. TC and EC interaction is described through the following reactions:

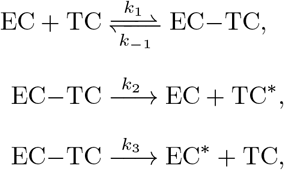

where EC – TC denotes conjugates of effector and tumor cells; and EC* and TC* are the inactivated effector and lithally-hit tumor cells, respectively. We define *E*, *T*, *C*, *E*\*, *T*\* as the number of EC, TC, EC–TC conjugates, EC*, and TC*, respectively. The non-negative kinetic parameters, *k*_1_, *k*_−1_, *k*_2_ and *k*_3_, describe the rates of the interactions. EC–TC conjugates can reversibly decompose without damaging the cells with the kinetic rate *k*_−1_. However, they can also irreversibly result in EC* or TC* with respective kinetic rates *k*_2_ and *k*_3_. The following system of nonlinear differential-algebraic equations describes those interactions, which is a slightly simplified version of the model of Kuznetsov [16].

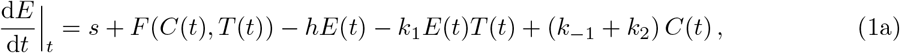

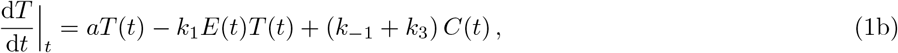

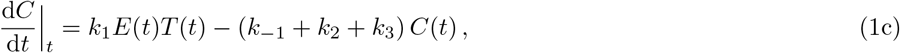

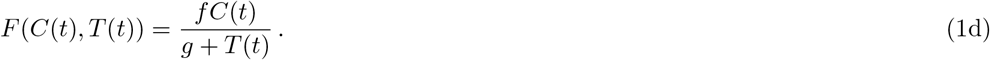

The equations describing the rate of change of *E*\* and *T*\* are not included in the system because they are irreversibly formed and thus have no effect on the other variables, and our target is to model *T* and *E* only. In [16], the model includes a sink term in the rate of change equation of *T* that represents TC growth limitation due to biological environment conditions. It considers, for example, resources competition like oxygen and substrates. We do not consider this term because growth limitations of even high initial *T* are associated with high rates of cytotoxic EC accumulation as well as the absence of their activity suppression by TC [39].

The rate of flow of mature EC to TC localization area is characterized by the generation term *s*. This rate is unaffected by the presence of TC. The destruction or migration of EC from the localization region of TC is represented by the elimination rate *h*. The model does not take into account any TC or EC–TC conjugates migration. Both multiplication and death of TC are included in parameter *a* that characterizes the maximum growth rate of TC population. The function *F* (*C*, *T*) represents the accumulation rate of the cytotoxic EC in the TC localization area due to tumor existence (stimulated accumulation), where *f* and *g* are positive constants. The EC accumulation, *F* (*C*, *T*), is due to signals, like released cytokines, generated by the EC in EC-TC conjugates. Thus this stimulated accumulation has some maximum value when *T* becomes large.

Following the suggestion of Kuznetsov [16], we consider a quasi-steady-state assumption for (1c), i.e., d_*C*_/d_*t*_|_*t*_ ≈ 0. Thus, *C* ≈ *KET*, where *K* = *k_l_*/ (*k*_2_ + *k*_3_ − *k*_−1_). As a result, (1a) and (1b) become:

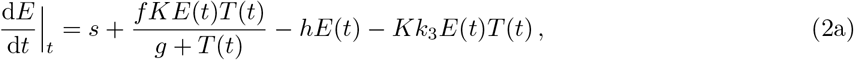

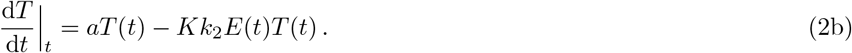

For further analysis and use of the model in parameter estimation and identifiability analysis, we use the same strategy of Kuznetsov [16] for non-dimensionalizing model equations. We non-dimensionalize (2a) and (2b) by considering concentration scales *E*_0_ = 10^7^ cells and *T*_0_ = 10^9^ cells for EC and TC, respectively [16]. We non-dimensionalize *t* by relating it to the deactivation rate of TC and introducing *τ* = *k*_2_*KT*_0_*t*/100. The final model formulation is:

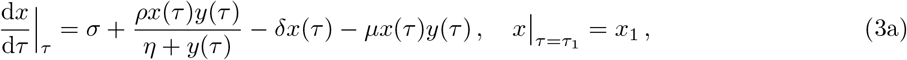

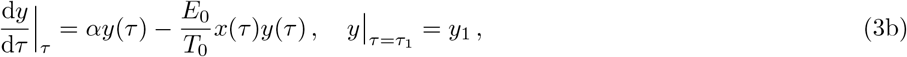

where

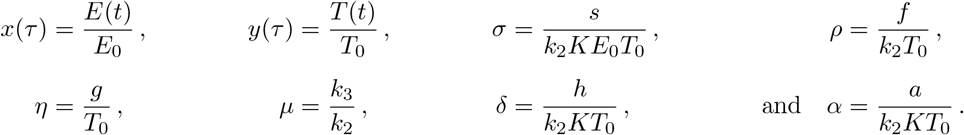

Thus, the final model is composed of two ordinary differential equations (ODEs) with two variables, *x* and *y*, with their respective initial values, *x*_1_ and *y*_1_, at the initial normalized time, *τ*_1_, and six unknown parameters, *σ*, *ρ*, *η*, *µ*, *δ*, and *α*.

### 3.2. Data Fitting and Parameter Estimation

To determine the parameter values for the nonlinear system (3a) and (3b) that best describe the experimental data, we conduct a regression analysis in the nonlinear least-squares sense by minimizing the sum of the squared residuals. The considered residuals are the differences between the measured values of tumor lesion longest diameters (converted to tumor number of cells as previously discussed) and the ones calculated from the model. In the present contribution, we do not seek global values of parameters, i.e, the same values for all patients. Instead, we solve the optimization problem for each patient individually to identify the parameter values that best describe the data of that patient. This constitutes a first step to check whether the model can describe the experimental data at all and analyze the ranges of parameter values. We also estimate the initial normalized value of EC number, *x*_1_, because it is unknown. In contrast, the initial normalized value of TC number, *y*_1_, is provided experimentally and does not need to be estimated. The nonlinear least square problem for each patient 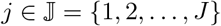 is expressed as follows:

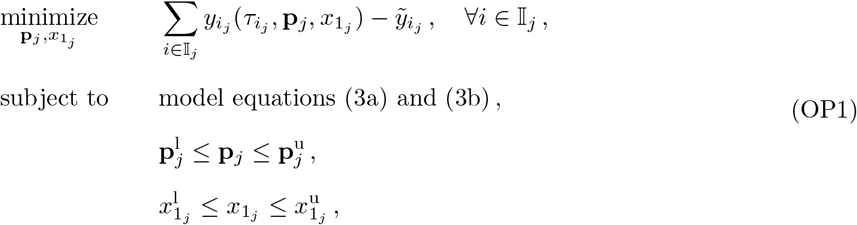

where *J* is the total number of patients considered. After data pre-processing, *J* = 210 patients. The index 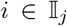 is for the observed experimental values, where 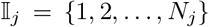 with *N*_*j*_ being the total number of observed values for patient *j*. The non-dimensionalized model-predicted and observed values of TC number at the normalized time 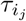 are 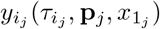 and 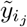, respectively. The initial value of the non-dimensionalized EC number at 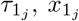, has lower and upper bounds 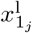 and 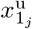, respectively. The vector **p**_*j*_ contains the non-dimensionalized parameter values of the model equations, (3a) and (3b), for patient *j*, **p**_*j*_ = [*σ_j_*, *µ_j_*, *δ_j_*, *α_j_*, *ρ_j_*, *η_j_*]. The lower and upper bounds of the components of **p**_*j*_ are the components of vectors 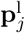 and 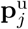, respectively. The decision variables of the optimization problem are thus the components of **p**_*j*_ and 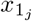. The dynamic optimization problem (OP1) is nonconvex and nonlinear, and can thus have multiple (suboptimal) local solutions. Hence, global optimization techniques are required to guarantee the global optimal solution, 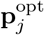 and 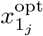.

### 3.3. Parameter Identifiability Analysis

Parameter identifiability analysis determines if model parameters can be uniquely estimated [40]. Different definitions of identifiability analysis are available in the literature. Miao et al. [41] reviewed several methods of parameter identifiability analysis for nonlinear ODE models and distinguished between different methodologies including structural and practical identifiability analyses. In the former analysis, one determines if a given structure of a model allows the realization of unique parameters when certain measured variables are provided [40]. However, it only provides necessary conditions for identifiability because it does not take into consideration parameters precision [42], [43]. In contrast, practical identifiability aims to predict confidence intervals for the estimated parameters [42], [44]. It can be conducted locally (in the neighborhood of the estimated parameter values) or globally over the entire range of values. We here carry out the latter analysis and evaluate it globally to improve the confidence in the parameter values that are determined by solving (OP1).

We conduct the global practical identifiability analysis by determining the smallest box that contains the so-called feasible parameter set 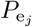 for each patient *j*, as suggested in [42]. This set includes all values of parameters in **p**_*j*_ for which the differences between model predictions 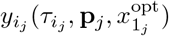 and optimal model predictions 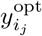 (determined by solving (OP1)) fall within certain defined bounds 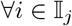, that is

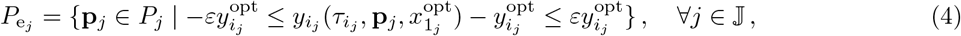

where 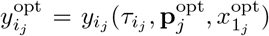, *ε* is the percentage of deviation, and *P_j_* is the set of parameter values in **p**_*j*_ bounded by the components of 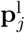 and 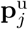 defined in (OP1). The set 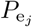 is depicted in dark gray as shown in Figure 2 for the case of a two-dimensional vector **p**_*j*_. We approximate the nonconvex set 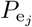 by a rectangular box (light gray color), whose edges are formed by the extreme values of the elements of **p**_*j*_ 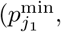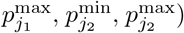. We determine these extreme values by solving a series of constrained dynamic optimization problems. For a vector **p**_*j*_ that consists of *K* parameters, the optimization problem for parameter number *k* is formulated 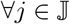, and 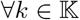 as follows [42], [45]:

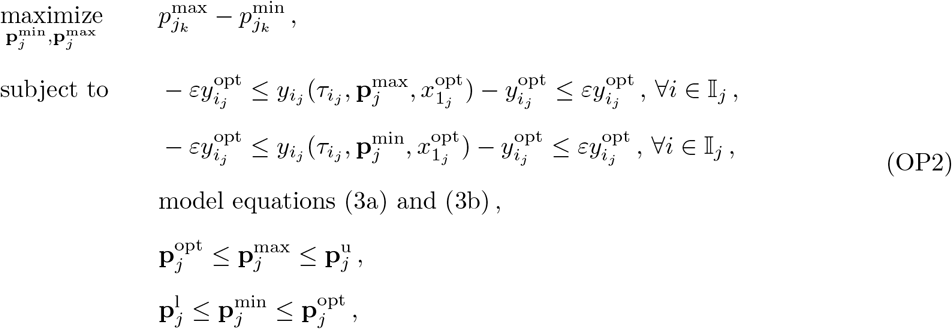

with 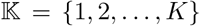 and *K* = 6. We set *ε* to 20%, which means that the differences between model predictions for 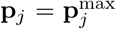 and 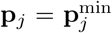, and optimal model predictions for 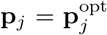 fall within 20% of the optimal prediction values 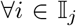. For each parameter *p_j_k* in **p**_*j*_, its minimum value 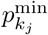 in 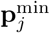 and its maximum value 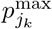 in 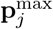 are determined by solving (OP2). Therefore, (OP2) is solved *K* times for each patient 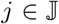. As a result, the approximation of 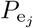 is determined, and its edges are the elements of 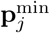 and 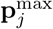. The set 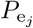 allows for the determination of confidence regions of the estimated parameter values in 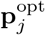. When 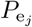 covers a large space in the direction of 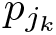, the estimated parameter 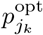 is not identifiable, in the sense that there is a large range of values for 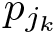 that could lead to a good fit. When the set covers a small space, 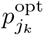 is identifiable because the parameter is determined to a sufficient accuracy. To decide on identifiability a certain threshold is thus needed. We here do not define a cutoff, we rather analyze the identifiability qualitatively.

**Figure 2:**
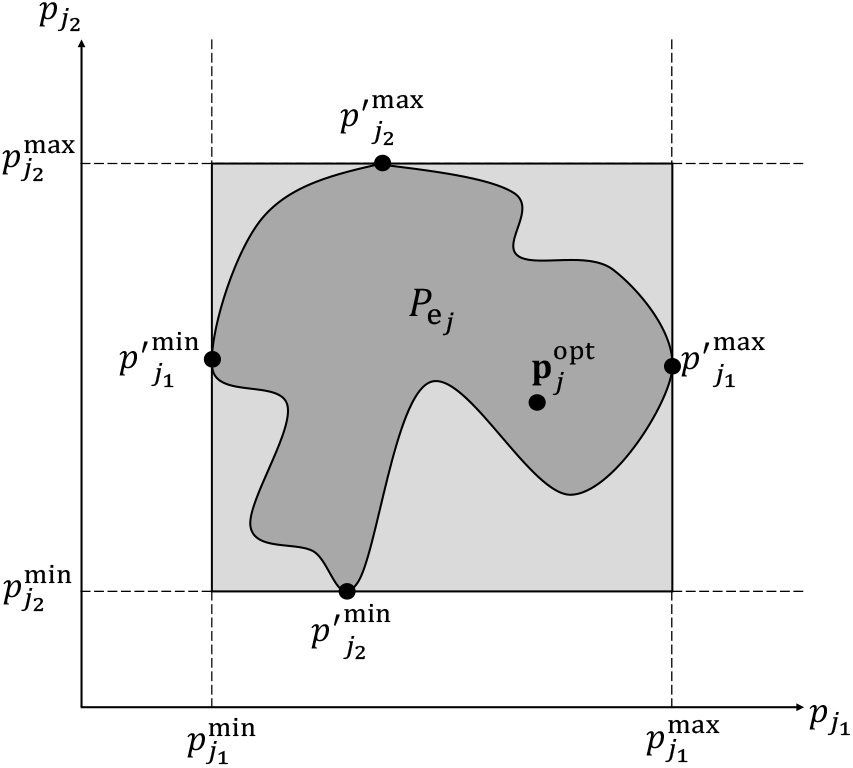
Illustration of parameter identifiability via global confidence intervals (based on [42]). The dark gray region indicates the feasible set 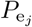 that includes all values of parameters in the two-dimensional vector **p**_*j*_ for which the differences between model predictions 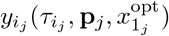 and optimal model predictions 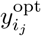 fall within certain defined bounds. The light gray region shows the rectangular box that conservatively approximates this feasible set, where the box edges are the extreme values of the elements of 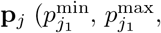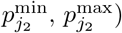.

### 3.4. Tumor Growth Prediction

Beyond being able to reproduce experimental data a posteriori, a more clinically relevant application of mathematical tumor modeling would be if the model was able to predict tumor growth. This could lead to model-based tumor treatment, as the supplied doses to patients could be adjusted in advance according to model predictions. In Subsection 3.2, we fitted the model to all experimental data points when estimating the parameters. In order to compare model extrapolation capabilities and future predictions to the clinical data, we now do not include the last *ζ* points of the data when fitting the model and solving (OP1). For each patient *j*, we solve an optimization problem similar to (OP1), but using only the data points in the set 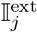 instead of 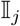, where 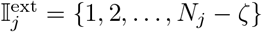. The optimal values of the decision variables obtained from this problem are called 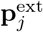 and 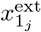. We then integrate the model equations for 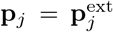 and 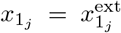 from 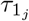 to 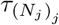, thus extrapolating beyond the data used for fitting. The results of the integrated 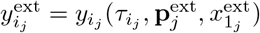 between 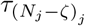 and 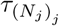 are the extrapolated part of the model that can be compared with the remaining *ζ* data points to gauge the extrapolation capabilities. We consider two model extrapolation cases in which *ζ* is equal to two and three.

Moreover, we formulate another optimization problem to investigate how far model extrapolation could deviate from the actual values. We aim to find two “extreme-case” lines that are designed to be as far away from each other at the final time point 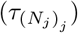 while both being within some *θ* tolerance of the found optimal fit 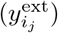 for the fitted time before extrapolation starts. For this, we solve the following optimization problem 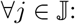:

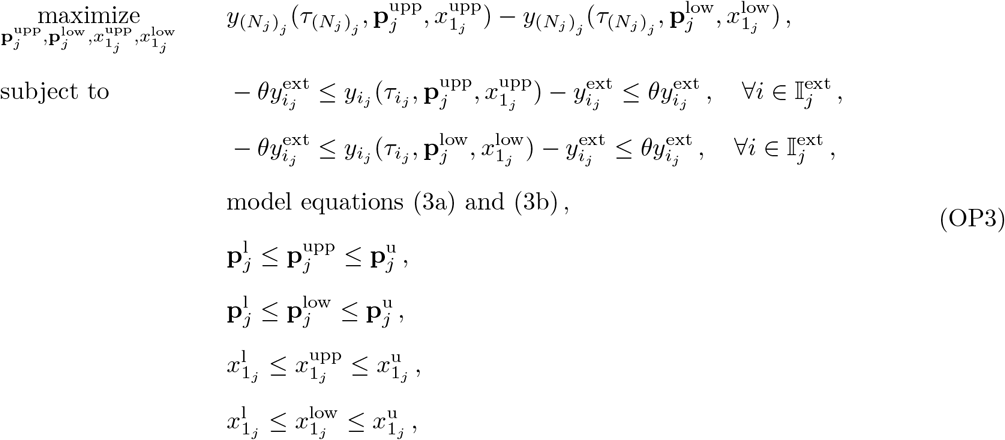

where 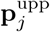 and 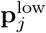 are the vectors that contain the parameter values for the upper and lower “extreme-case” model extrapolation deviations, respectively. The initial values of the non-dimensionalized EC number at 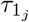 for these upper and lower deviation cases are 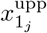 and 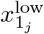, respectively. We set *θ* to 10%. By integrating (3a) and (3b) from 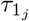 to 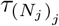, we get 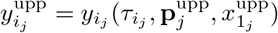 and 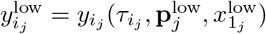, which allow the comparison of these two “extreme-case” extrapolation deviations with the remaining *ζ* data points after 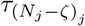 (start of extrapolation).

### 3.5. Implementation

We implement the model, (3a) and (3b), in MATLAB R2019b [46]. All optimization problems are solved in the MATLAB version of the global optimization toolbox MEIGO using the enhanced scatter search metaheuristic (eSS) method [47]. The eSS is stochastic and employs some elements of the scatter search and path re-linking methodologies [48]. Thus, the solution depends on the initial conditions and the global optimum is not guaranteed. We set the maximum number of function evaluations, the maximum CPU time and the maximum absolute violation of the constraints to 10^5^, 100 s and 10^−5^, respectively. We use a 50% probability of biasing the search towards bounds and the dynamic hill climbing (DHC) [49] as a local search method. For all aforementioned optimization problems, we set 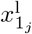 and 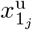 to 10^−2^ and 10^2^, respectively. All elements of 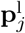 and 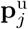 are set to 10^−2^ and 10^2^, respectively.

The ODE (3a) and (3b) are solved using the variable-step, variable-order (VSVO) solver based on the numerical differentiation formulas (NDFs) of orders one to five (ode15s) [50]. We set the relative and absolute error tolerances to 10^−3^ and 10^−6^, respectively. The solution refinement factor is one and the maximum step size is 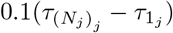.

## 4. Results and Discussion

We now show the results of estimation, identifiability, and predictions. We show the results of six selected patients in details and give performance measures for all 210 patients. We selected those six patients in a way to provide the different profiles of TC dynamics. Data fitting and growth prediction results of all 210 patients are provided in the supplementary material.

### 4.1. Data Fitting and Parameter Estimation

We fitted the parameters of a modified Kuznetsov model on a dataset of solid tumors in human patients under immunotherapy treatment. The model predicted the different tumor growth profiles represented by a selection of representative patients, as well as in the total (unselected) cohort. As shown in Figure 3 and Table 2, model prediction and experimental data profiles are qualitatively and quantitatively very close for these patients. The mean absolute error (MAE), the root-mean-square error (RMSE), and the coefficient of determination (*R*^2^) for the selected six patients are given in Table 2. We found that the model gave very high goodness of fit as measured by *R*^2^. Across 210 patients in all studies, an average *R*^2^ of 0.784 was achieved. Moreover, Table 4 provides the number of patients and the *R*^2^ values per study and per arm. Study 1 and Study 5 have higher *R*^2^ than the remaining studies. Arm 1 has the highest *R*^2^ in all studies except for Study 2. Although direct comparison of this performance with the previous work in [30] is not possible, comparing the MAE of the selected six patients with the reported average MAE in the previous study indicates good fitting performance of the developed Kuznetsov model. Furthermore, by analyzing the goodness of fit in individual patients, we found that the modified Kuznetsov model was able to fit clinically interesting patterns. In particular, the modified Kuznetsov model was able to predict relapse after initial tumor response (patient #207 in Figure 3) and other types of fluctuating behavior, solving a key limitation of previously used simpler models as in [30].

**Table 2:** Quantification of the goodness of fit of the model as shown in Figure 3. We compare the values of all data points to model results when calculating MAE, RMSE and *R*^2^.

| Patient # | MAE | RMSE | $R^2$ |
| --- | --- | --- | --- |
| 22 | 0.178 | 0.226 | 0.931 |
| 83 | 0.065 | 0.103 | 0.996 |
| 163 | 0.084 | 0.108 | 0.963 |
| 186 | 1.171 | 1.993 | 0.949 |
| 203 | 1.638 | 2.039 | 0.974 |
| 207 | 0.115 | 0.154 | 0.987 |
| All Patients (1 → 210) | — | — | 0.784 |

**Table 3:**
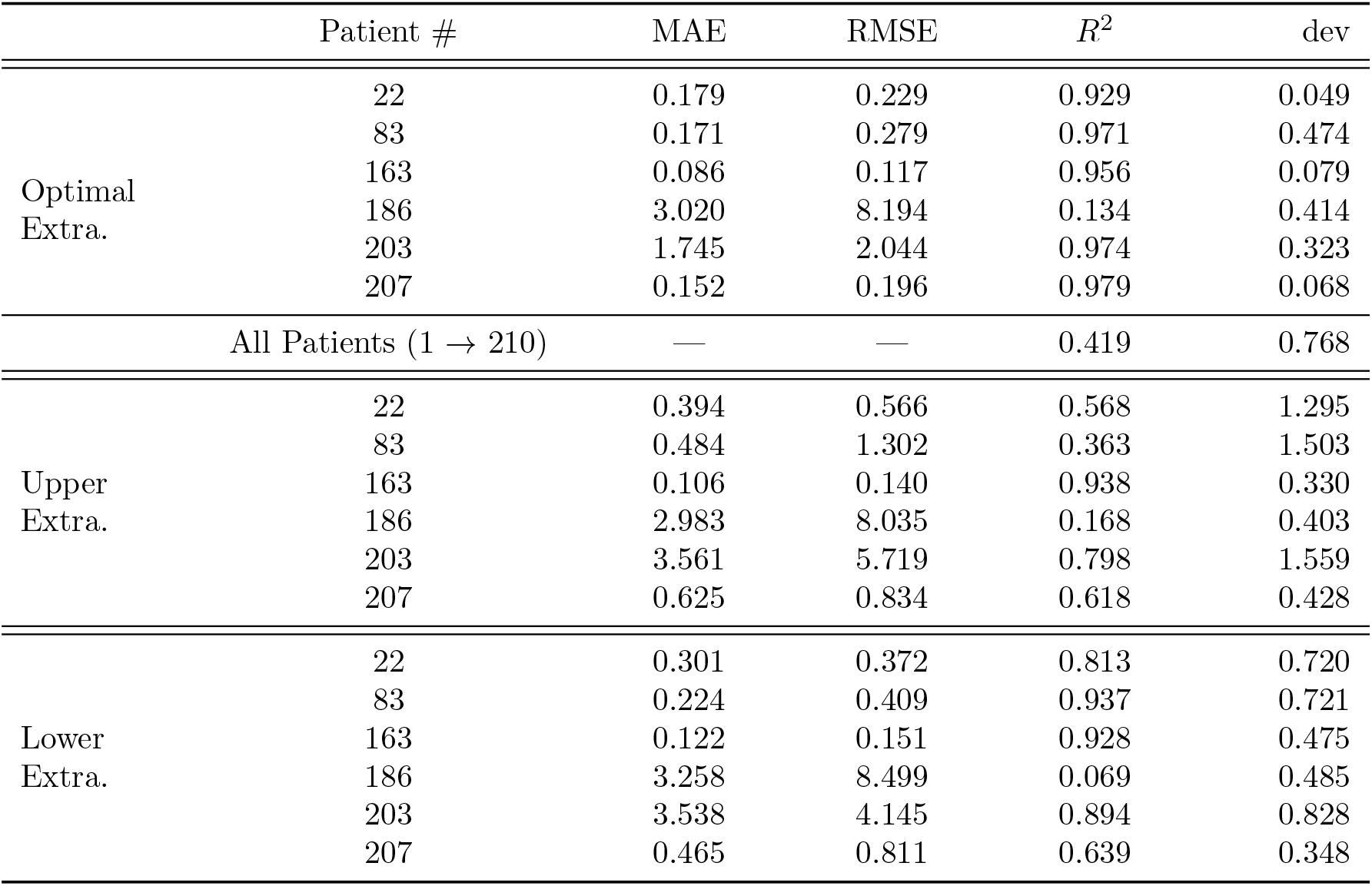
Quantification of the goodness of fit and prediction of the model as shown in Figure 6. Here the last two data points are not considered for parameter estimation. We compare the values of all data points to model results when calculating the errors and *R*^2^. The average deviation of model prediction results from the measured values during extrapolation time (shaded region in Figure 6) is represented by dev. The Optimal Extra. sub-table shows the values for the optimal model prediction results (the solid (black) line in Figure 6). The Upper Extra. sub-table relates values to the upper “extreme-case” extrapolation deviation results (the dash-dotted (blue) line in Figure 6). The Lower Extra. sub-table relates values to the lower “extreme-case” extrapolation deviation results (the dashed (magenta) line in Figure 6).

**Table 4:** Quantification of the goodness of fit and extrapolation of the model per the conducted studies and arms. We compare the values of all data points to model results when calculating the *R*^2^. For model extrapolation, the average deviation of model prediction results from the measured values during extrapolation time (shaded region in Figure 6) is represented by dev. A detailed description of the included studies is available in [30]. Briefly, study arms with “Atezolizumab” or “MPDL” are immunotherapy. “Docetaxel” is the main chemotherapy in these trials.

| Study No. (Name) | Arm No. (Name) | No. of Patients | Data Fitting | Extrapolation |  |
| --- | --- | --- | --- | --- | --- |
| | | | $R^2$ | $R^2$ | dev |
| 1 (FIR) | 1 (MPDL3280A-1) | 3 | 0.869 | 0.582 | 1.256 |
|  | 2 (MPDL3280A-2) | 11 | 0.847 | 0.535 | 0.406 |
|  | 3 (MPDL3280A-3) | 0 | — | — | — |
|  | Total | 14 | 0.852 | 0.545 | 0.588 |
| 2 (POPLAR) | 1 (Atezolizumab) | 6 | 0.581 | 0.241 | 0.943 |
|  | 2 (Docetaxel) | 25 | 0.747 | 0.474 | 0.761 |
|  | Total | 31 | 0.715 | 0.429 | 0.796 |
| 3 (BIRCH) | 1 (MPDL3280A-1a) | 2 | 0.964 | 0.980 | 1.983 |
|  | 2 (MPDL3280A-2a) | 6 | 0.587 | -0.077 | 1.143 |
|  | 3 (MPDL3280A-3a) | 2 | 0.243 | -0.036 | 0.317 |
|  | 4 (MPDL3280A-1b) | 8 | 0.864 | 0.522 | 0.525 |
|  | 5 (MPDL3280A-2b) | 3 | 0.749 | 0.226 | 0.394 |
|  | 6 (MPDL3280A-3b) | 9 | 0.884 | 0.575 | 0.670 |
|  | Total | 30 | 0.769 | 0.382 | 0.762 |
| 4 (OAK) | 1 (Atezolizumab) | 29 | 0.811 | 0.218 | 0.393 |
|  | 2 (Docetaxel) | 82 | 0.783 | 0.455 | 0.869 |
|  | Total | 111 | 0.790 | 0.393 | 0.745 |
| 5 (IMvigor 210) | 1 (Atezolizumab) | 24 | 0.825 | 0.501 | 0.953 |
| Total |  | 210 | 0.784 | 0.419 | 0.768 |

**Figure 3:**
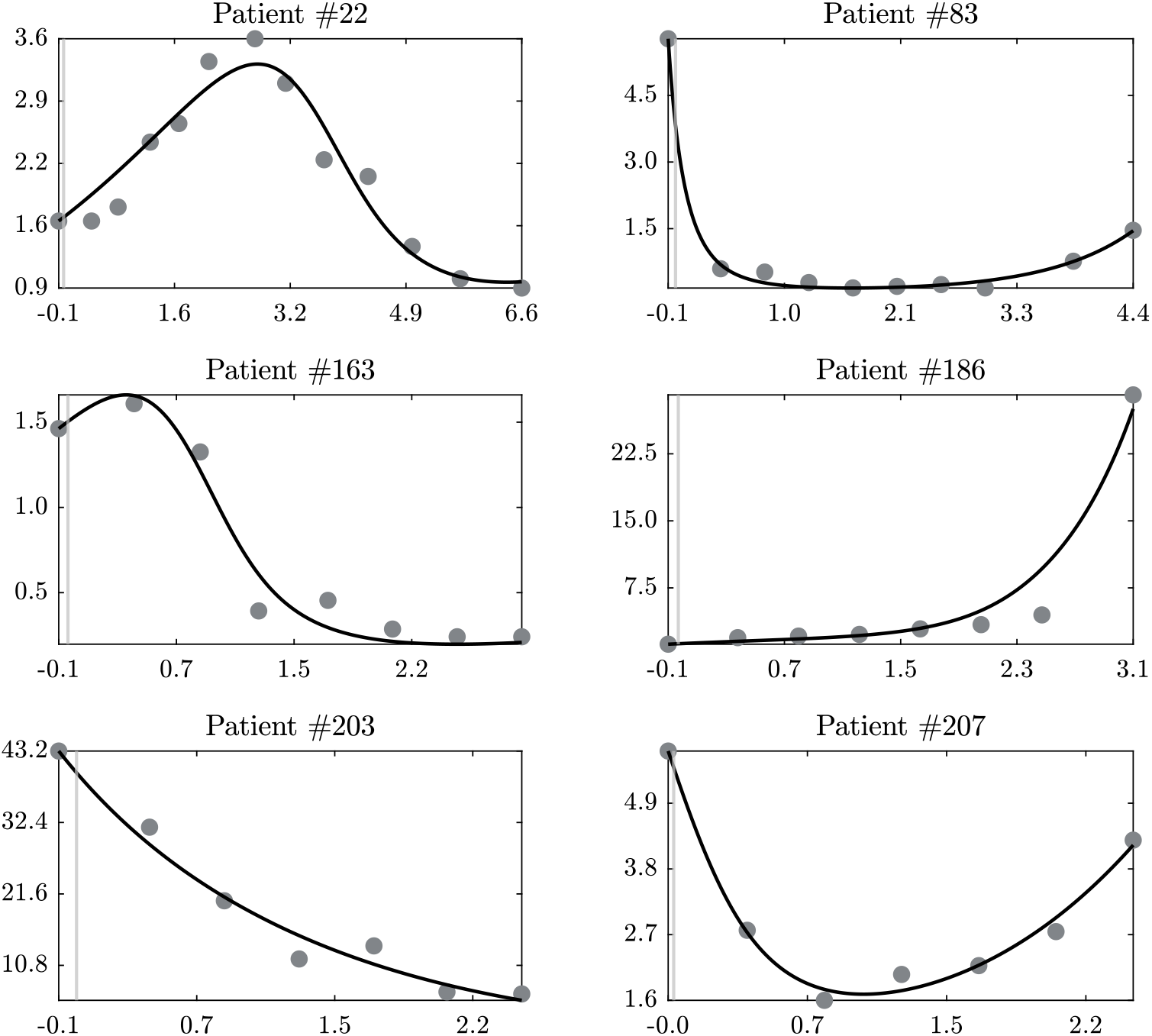
Data fitting results of TC number of the selected patients. The solid (black) line shows model results, where all data points are used when estimating the parameters. The points represent the measured data. Ordinates: normalized number of tumor cells. Abscissas: normalized treatment time, where negative values indicate time before the start of treatment. The model can fit experimental data with different qualitative trends (e.g., up, down and “U”-curve).

We fitted the parameters of the modified Kuznetsov model to a large clinical dataset obtained from five clinical trials. In general, the distributions of the resulting parameter values were similar between the studies (Figure 4). These ranges can be useful for further studies because they enable other researchers to determine plausible ranges and optimal boundaries when fitting the same model to other datasets, thereby simplifying the optimization procedure. A global parameter estimation, however, was not performed in this study and could be attempted in future studies.

**Figure 4:**
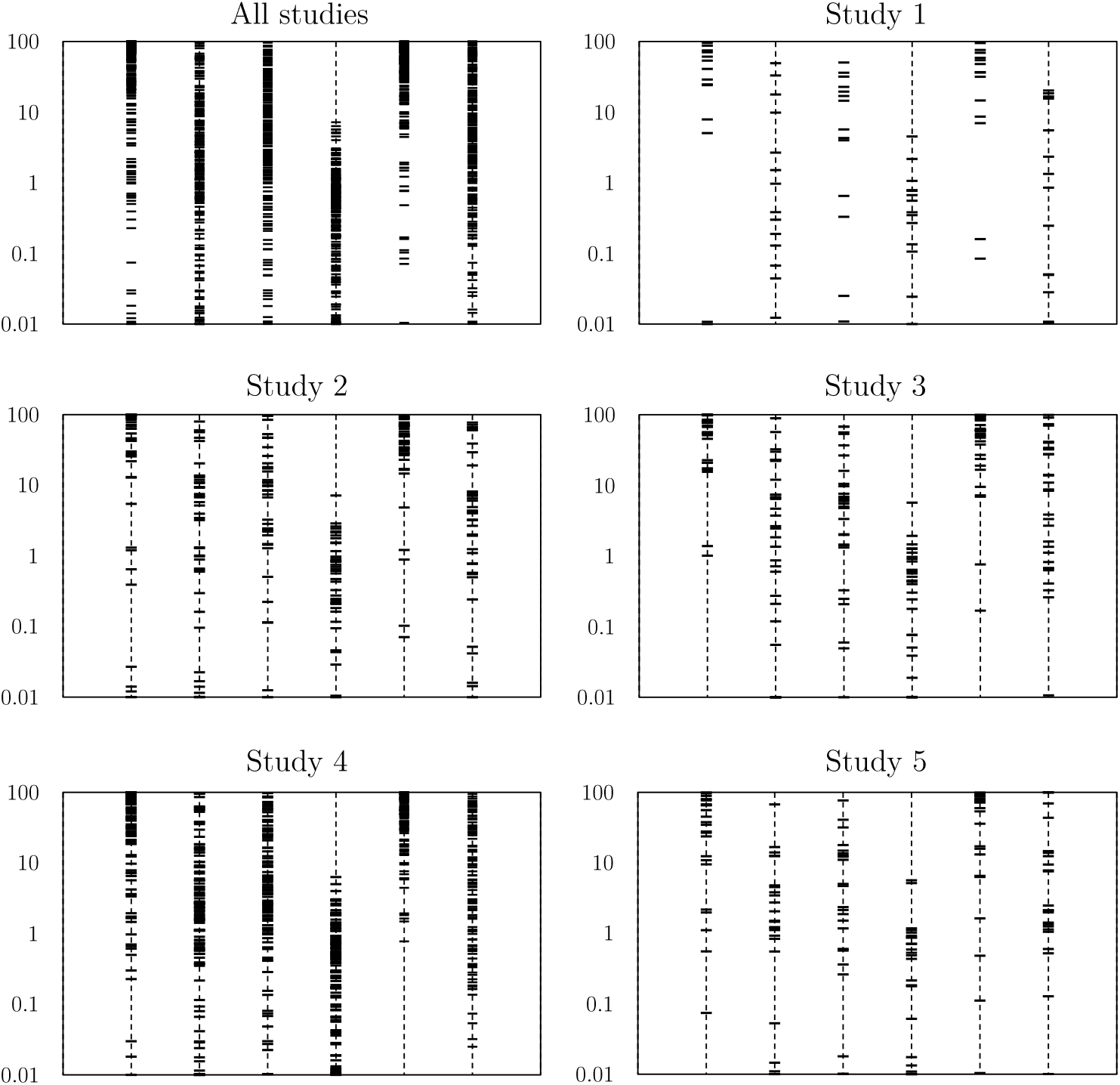
Estimated values of the parameters of (3a) and (3b) model for all 210 patients (after data pre-processing) individually. Ordinates: parameter values. Abscissas: non-dimensionalized model parameters, placed in the following order from left to right: *σ*, *µ*, *δ*, *α*, *ρ*, *η*. The values are scattered all over the bounds’ ranges, but *α* values, which have a maximum of 6.331. Moreover, the distribution densities of parameter values are very close to each other among the studies.

Figure 4 shows the estimated values of model parameters of all 210 patients. All parameter values vary per patient. The values are distributed all over the bounds, except for *α*, which has a maximum of 6.331. The parameter *α* is the normalized parameter for *a*, which represents the maximum growth rate of TC population. In addition, most of the values of *µ* and *ρ* are close to the upper bound. Although parameter values are quite distributed between the bounds, the aforementioned findings can help in narrowing the expected ranges of values of parameters when global parameter estimation is targeted. Moreover, the distribution of parameter values is compared among the considered five studies. As we can see in Figure 4, the distribution densities between the bounds of parameter values are the same for all studies.

### 4.2. Parameter Identifiability Analysis

Figure 5 provides the identifiability analysis results of the estimated model parameter values of the selected patients. Depending on the patient and the parameter, the estimated parameter values can be unique or take other values. For patient #22, the approximated feasible parameter set covers a small space in all parameters directions. Their estimated parameter values are close to being unique and thus identifiable. On the other hand, the approximated feasible parameter set for patient #203 covers a large space in all parameter directions. Therefore, the found values are not unique and other combinations of values could lead to good fits and predictions. For the remaining selected patients, the feasible parameter sets can be small or large depending on the patient and the parameter.

**Figure 5:**
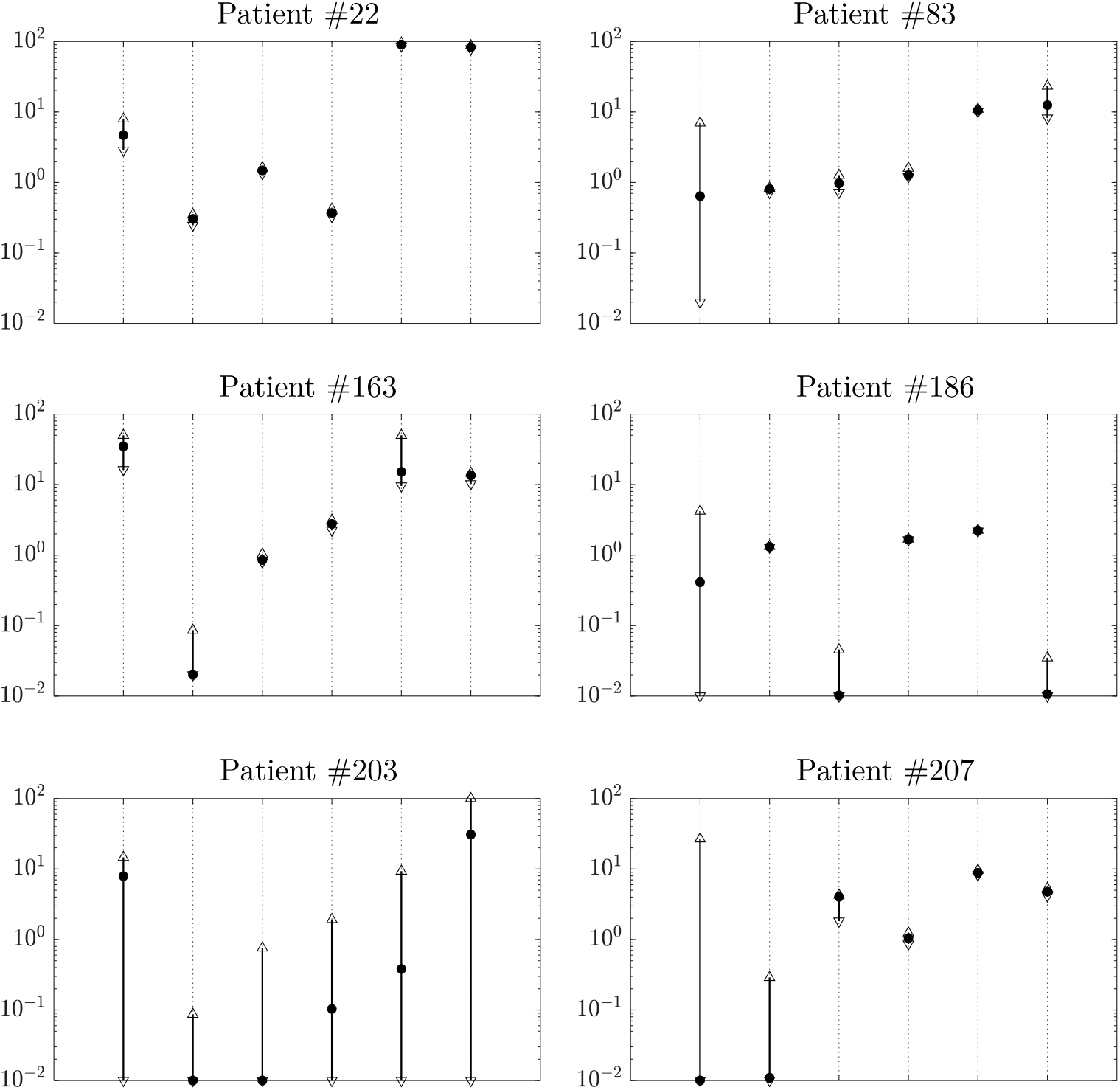
Identifiability analysis results of estimated values of model parameters of the selected patients. Ordinates: parameter values. Abscissas: non-dimensionalized model parameters, placed in the following order from left to right: *σ*, *µ*, *δ*, *α*, *ρ*, *η*. The points are the estimated values. The arrowheads are the maximum and minimum values found by the analysis. The results show that there are several combinations of parameter values at which the model can fit the experimental data.

When the target is to estimate parameter values of the tumor dynamics model of a certain patient, identifiable values are what we need. However, when the aim is to find global parameters values for all patients, large spaces of the feasible parameter sets can be desirable for finding those global values, in which they are independent of a considered patient. In summary, the results of Figure 5 show that for several combinations of parameter values, good data fitting and model predictions can be achieved. This opens the potential for global parameter estimation, in which the estimated values of model parameters are the same for all patients.

### 4.3. Tumor Growth Prediction

In clinical decision making, a possible role of mechanistic models is forecasting tumor growth during treatment, potentially enabling physicians to adjust the treatment strategy earlier. We found that the modified Kuznetsov model indeed was able to extrapolate beyond the initial time points when the last two or three data points are not included when fitting the model. The solid black lines in Figure 6 show model extrapolation results of the selected patients when the last two data points are not considered for fitting. For the six patients, the model quantitatively forecasts tumor dynamics, except for patient #186. The last data point of patient #186 is almost impossible to forecast because it suddenly shifts upward after a mild and constant increase of tumor growth. However, the model can still qualitatively predict the growth. For the other patients, the predictions are very close to the experimental data. In Table 3, the first sub-table “optimal Extra.” provides the MAE, RMSE, and *R*^2^ of the selected patients, as well as the average deviation for the extrapolated part, represented as “dev”. The values of *R*^2^ for the six patients are close to one except for patient #186 due to the aforementioned explanation. Compared to the previous work in [30], model extrapolation results have here higher *R*^2^ values, specifically, an *R*^2^ of 0.979 was reached for patient #207. The average values of *R*^2^ and dev of all 210 patients are also provided in the table. Model extrapolation results when omitting the last two data points for all 210 patients are provided in the supplementary material. Also, we provide model extrapolation results while omitting the last three data points for the selected six patients in the supplementary material. For the latter results, the model could also predict the growth very well.

**Figure 6:**
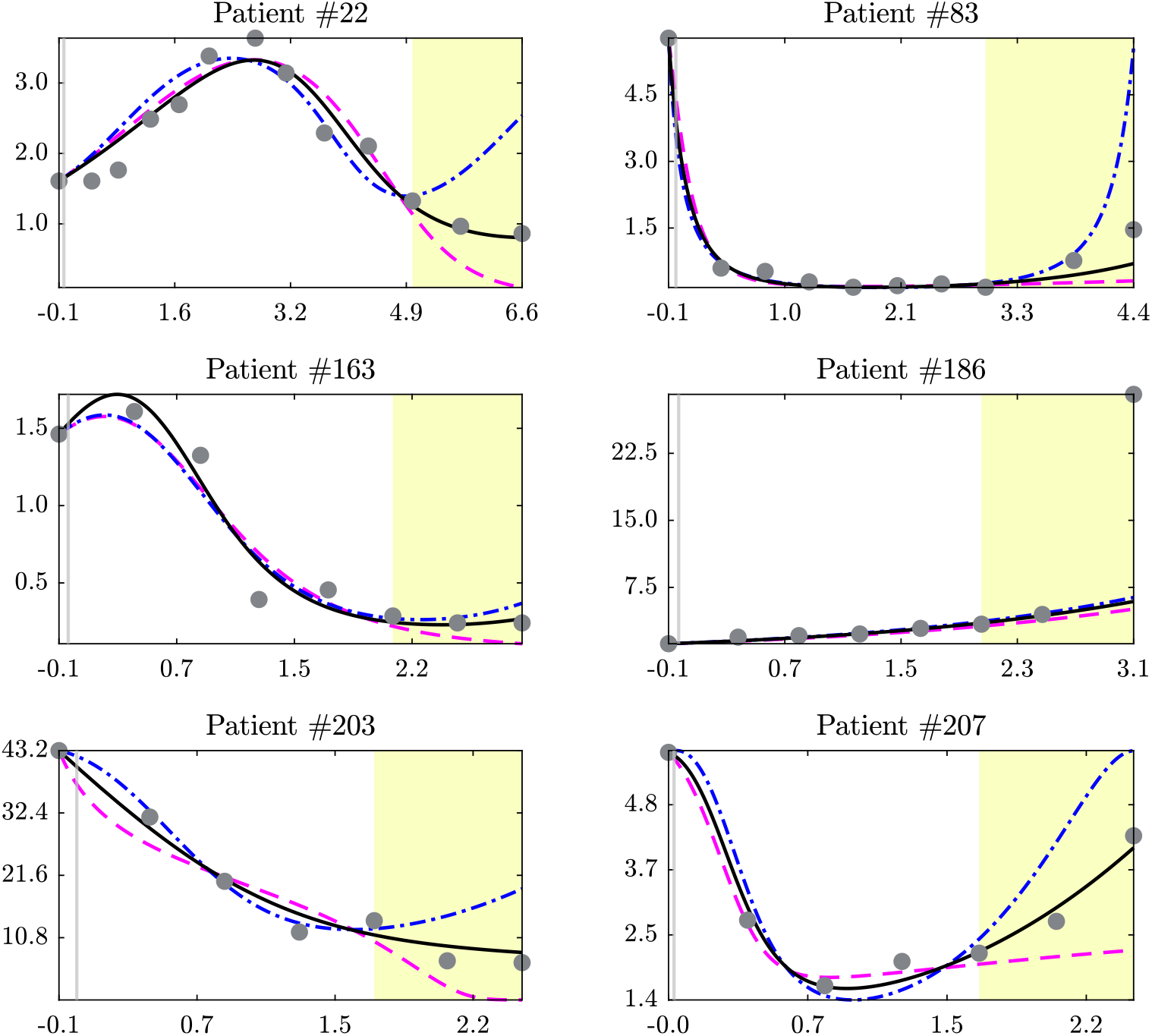
Model extrapolation results of the selected patients. The solid (black) line represents the optimal model prediction. The dash-dotted (blue) and the dashed (magenta) lines show the upper and the lower “extreme-case” model extrapolation deviation results, respectively. Here the last two data points are not considered for parameter estimation. The points show the measured data. Shaded areas highlight regions of model extrapolation. Ordinates: normalized number of tumor cells. Abscissas: normalized treatment time, negative values indicate time before the start of treatment. The model is capable of forecasting tumor dynamics qualitatively and sometimes quantitatively.

In addition, Table 4 provides the *R*^2^ and dev values per study and per arm for model extrapolation when omitting the last two data points. In general, and as expected, the calculated average *R*^2^ for the extrapolation experiment is lower than the *R*^2^ when using all the data points in all the studies. Similar to data fitting, Study 1 and Study 5 have higher *R*^2^ than the remaining studies. In Study 3, Arm 2 and Arm 3 have the lowest *R*^2^. Interestingly, in Study 2 and Study 4, the extrapolation performance of the model is markedly better for the Docetaxel group (Arm 2) than for the Atezolizumab group (Arm 1). This indicates that patients receiving Docetaxel have a more predictable response trajectory compared to patients receiving immunotherapy in whom unexpected patterns of tumor response can occur even at later time points.

Moreover, Figure 6 shows how far model extrapolation could deviate from the actual values. The results of the two “extreme-case” (dash-dotted blue and the dashed magenta) lines are designed to be as far away from each other at the final time point 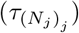 while both being within some 10% tolerance of the found optimal fit for the fitted time before extrapolation starts. For some patients, the “extreme-case” lines can significantly deviate from the actual values, especially for the upper case (dash-dotted blue lines), as for patients #22 and #83. In contrast, the “extreme-case” lines for patients #163 and #186 are very close to the optimal extrapolated ones. Sub-tables “Upper Extra.” and “Upper Extra.” in Table 3 provide the MAE, RMSE, *R*^2^ and dev for the selected patients for the two “extreme-case” extrapolations.

To sum up, the model can forecast tumor dynamics of the patients and the “extreme-case” extrapolation scenarios were conducted to check the worst model predictions. For some patients the “extreme-case” extrapolation results deviate from the actual values, for others they remain close to the optimal extrapolation results.

## 5. Conclusion

In this study, we show that quantitative mathematical models can be used to describe and forecast the behavior of cancer. Previous studies have used the same datasets to fit very simple ODE models to the tumor volume measurements over time [30]. However, in Ghaffari Laleh et al. [30], it was observed that all established ODE models were not able to fit “U”-shaped trajectories well. In clinical terms, patients who relapsed after an initial response, or patients who showed a delayed response, were not adequately represented in these previous models. Compared to this, the present study evaluates a more complex model which has the benefit of being able to describe a larger variety of real-world time series. This specific model is a slight simplification of the Kuznetsov model [16], which has not been linked with or validated in large amounts of quantitative real-world human data. Specifically, it could quantitatively fit the Kuznetsov model to a large dataset of 1472 patients. Data are collected from patients undergoing immunotherapy or chemotherapy treatments.

In the parameter estimation for each patient, we found that some parameters for some patients are not unique (identifiability analysis). This means that many combinations of parameter values could lead to good fitting and predictions. This opens the potential for global parameter estimation, in which parameter values are the same for all patients. Still, the model could predict tumor growth (2-3 omitted measurements) which could indicate practical usefulness as a predictive biomarker. Specifically, the model fitting and prediction could potentially describe and forecast the behavior of cancer, improve the understanding of underlying biological mechanisms, and provide model approaches for cancer treatments. Future studies should attempt a global parameter estimation, use deterministic and less computationally demanding solution methods. Another possibility is the reduction of the number of model parameters in order to have less number of parameters to estimate and thus enhance the possibility to reach global values.

The model complexity is mostly limited by the availability of suitable data. Therefore, if more data become available, it would be possible to include other terms in the model, e.g., explicitly representing different types of immune cell populations or multiple cancer cell clones.

## 6. Additional Information

### 6.1. Author Contributions

MEW performed the numerical experiments; NGL and JNK selected the publicly available data and provided clinical insight; MEW, FJ, DB, CK, AM analyzed the data; MEW, NGL and JNK wrote the initial version of the manuscript; all authors contributed to the interpretation of the results; all authors edited the manuscript and made the decision to submit for publication.

### 6.2. Competing Interests

None.

## 6.3. Acknowledgments

None.

## Notes

https://www.biorxiv.org/content/10.1101/2021.10.23.465549v2.supplementary-material

### Competing Interest Statement

The authors have declared no competing interest.

