## Supplementary Information for "Can the Kuznetsov Model Replicate and Predict Cancer Growth in Humans?"

### Supplementary Material for: Can the Kuznetsov Model Replicate and Predict Cancer Growth in Humans?

Estimated values of the model parameters of the 111 patients from study 4 (after data pre-treatment), shown per each arm in the study. Ordinates: parameter values. Abscissas: non-dimensionalized model parameters, placed in the following order from left to right: $\sigma$, $\mu$, $\delta$, $\alpha$, $\rho$, $\eta$. The values are scattered all over the bounds' ranges, but $\alpha$ values, which have a certain maximum. Moreover, the distribution densities of parameter values are very close to each other between the two arms.

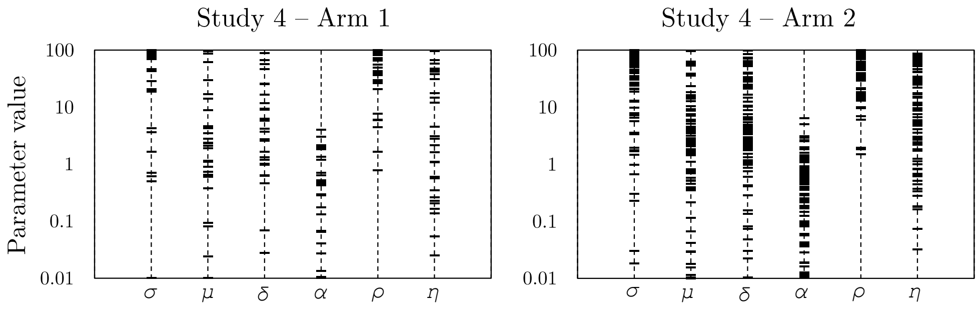

Model extrapolation results of the selected patients. The solid-line curves and the points represent model results and measured data, respectively. The last three data points are not used for parameters estimation. Ordinates: normalized number of tumor cells. Abscissas: normalized treatment time, negative values indicate time before the start of treatment. The model is capable of forecasting tumor dynamics qualitatively and sometimes quantitatively.

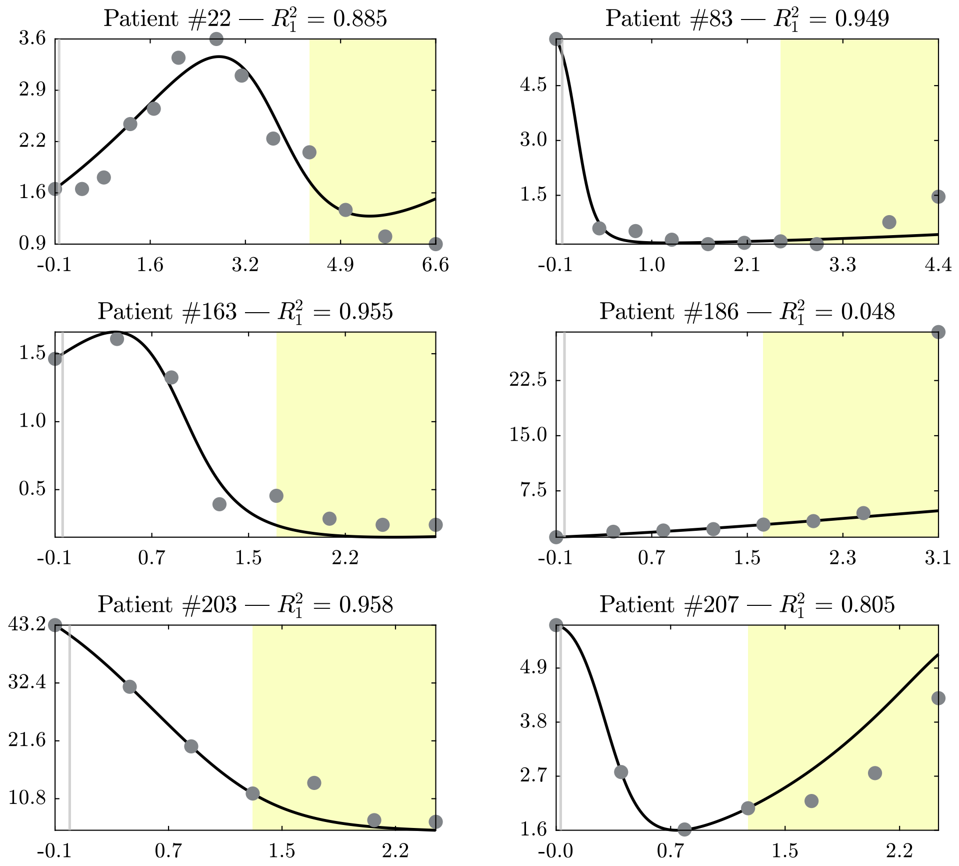

Data fitting results of TC number of all 210 patients. The solid (black) line shows model results, where all data points are used when estimating the parameters. The points represent the measured data. Ordinates: normalized number of tumor cells. Abscissas: normalized treatment time, where negative values indicate time before the start of treatment. The model can fit experimental data with different qualitative trends (e.g., up, down and “U”-curve).

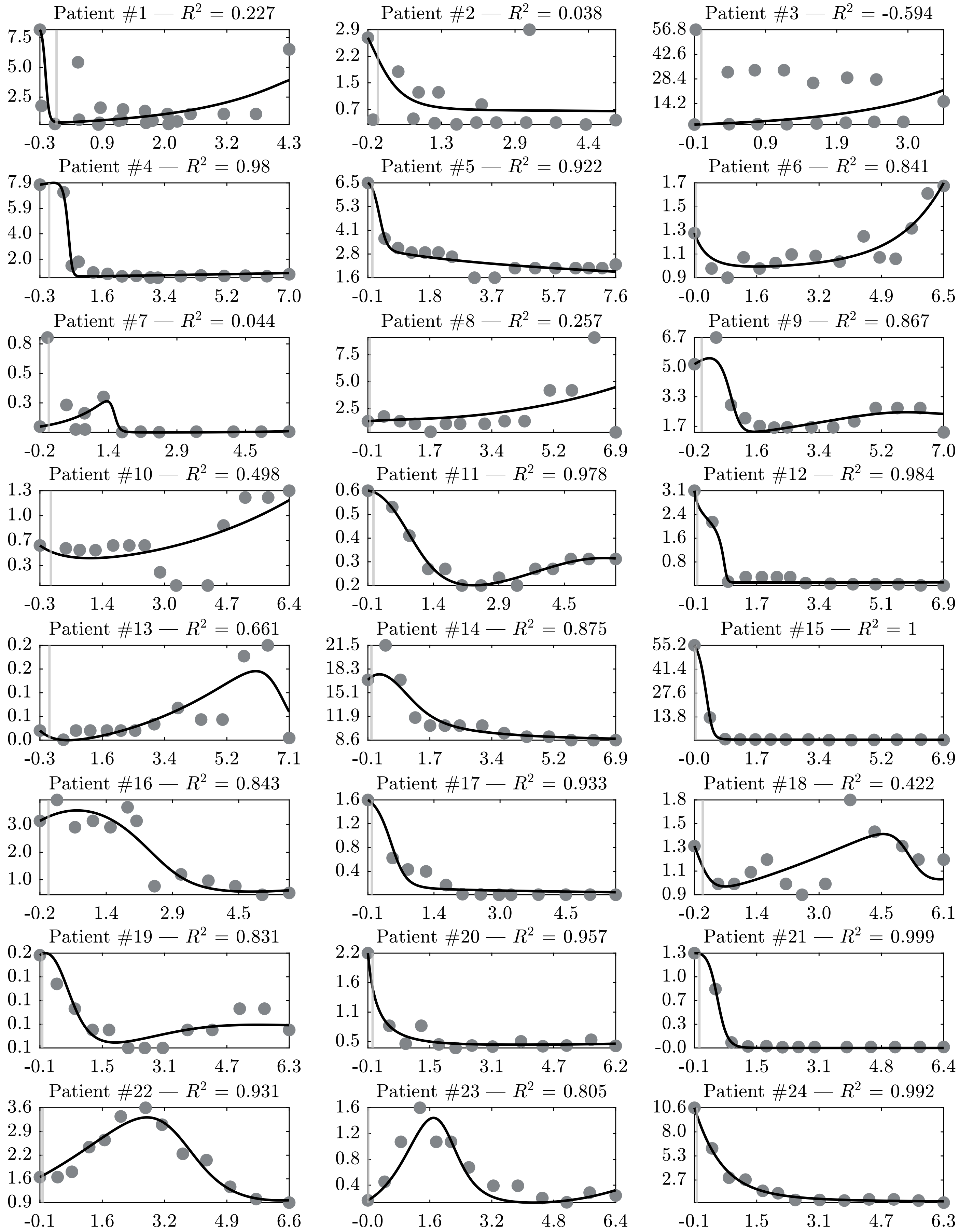

Patient #1 — $R^2$ = 0.227
Patient #2 — $R^2$ = 0.038
Patient #3 — $R^2$ = -0.594
Patient #4 — $R^2$ = 0.98
Patient #5 — $R^2$ = 0.922
Patient #6 — $R^2$ = 0.841
Patient #7 — $R^2$ = 0.044
Patient #8 — $R^2$ = 0.257
Patient #9 — $R^2$ = 0.867
Patient #10 — $R^2$ = 0.498
Patient #11 — $R^2$ = 0.978
Patient #12 — $R^2$ = 0.984
Patient #13 — $R^2$ = 0.661
Patient #14 — $R^2$ = 0.875
Patient #15 — $R^2$ = 1
Patient #16 — $R^2$ = 0.843
Patient #17 — $R^2$ = 0.933
Patient #18 — $R^2$ = 0.422
Patient #19 — $R^2$ = 0.831
Patient #20 — $R^2$ = 0.957
Patient #21 — $R^2$ = 0.999
Patient #22 — $R^2$ = 0.931
Patient #23 — $R^2$ = 0.805
Patient #24 — $R^2$ = 0.992

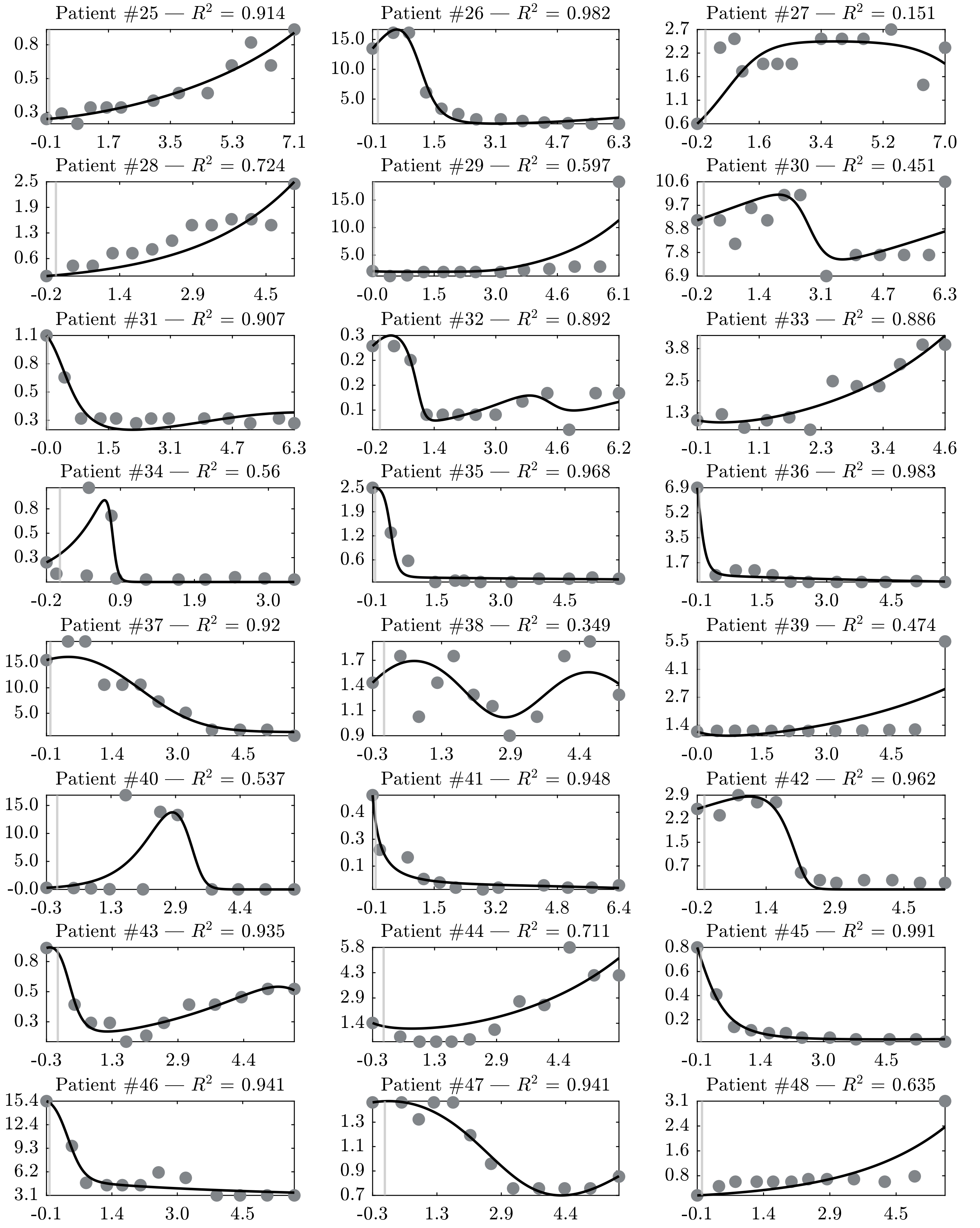

Patient #25 — $R^2$ = 0.914
0.8
0.5
0.3
-0.1
1.7
3.5
5.3
7.1
Patient #26 — $R^2$ = 0.982
15.0
10.0
5.0
-0.1
1.5
3.1
4.7
6.3
Patient #27 — $R^2$ = 0.151
2.7
2.2
1.6
1.1
0.6
-0.2
1.6
3.4
5.2
7.0
Patient #28 — $R^2$ = 0.724
2.5
1.9
1.3
0.6
-0.2
1.4
2.9
4.5
Patient #29 — $R^2$ = 0.597
15.0
10.0
5.0
-0.0
1.5
3.0
4.6
6.1
Patient #30 — $R^2$ = 0.451
10.6
9.7
8.8
7.8
6.9
-0.2
1.4
3.1
4.7
6.3
Patient #31 — $R^2$ = 0.907
1.1
0.8
0.5
0.3
-0.0
1.5
3.1
4.7
6.3
Patient #32 — $R^2$ = 0.892
0.3
0.2
0.2
0.1
-0.2
1.4
3.0
4.6
6.2
Patient #33 — $R^2$ = 0.886
3.8
2.5
1.3
-0.1
1.1
2.3
3.4
4.6
Patient #34 — $R^2$ = 0.56
0.8
0.5
0.3
-0.2
0.9
1.9
3.0
Patient #35 — $R^2$ = 0.968
2.5
1.9
1.2
0.6
-0.1
1.5
3.0
4.5
Patient #36 — $R^2$ = 0.983
6.9
5.2
3.5
1.7
-0.1
1.5
3.0
4.5
Patient #37 — $R^2$ = 0.92
15.0
10.0
5.0
-0.1
1.4
3.0
4.5
Patient #38 — $R^2$ = 0.349
1.7
1.4
1.1
0.9
-0.3
1.3
2.9
4.4
Patient #39 — $R^2$ = 0.474
5.5
4.1
2.7
1.4
-0.0
1.5
3.0
4.5
Patient #40 — $R^2$ = 0.537
15.0
10.0
5.0
-0.0
-0.3
1.3
2.9
4.4
Patient #41 — $R^2$ = 0.948
0.4
0.3
0.1
-0.1
1.5
3.2
4.8
6.4
Patient #42 — $R^2$ = 0.962
2.9
2.2
1.5
0.7
-0.2
1.4
2.9
4.5
Patient #43 — $R^2$ = 0.935
0.8
0.5
0.3
-0.3
1.3
2.9
4.4
Patient #44 — $R^2$ = 0.711
5.8
4.3
2.9
1.4
-0.3
1.3
2.9
4.4
Patient #45 — $R^2$ = 0.991
0.8
0.6
0.4
0.2
-0.1
1.4
3.0
4.5
Patient #46 — $R^2$ = 0.941
15.4
12.4
9.3
6.2
3.1
-0.1
1.4
3.0
4.5
Patient #47 — $R^2$ = 0.941
1.3
1.1
0.9
0.7
-0.3
1.3
2.9
4.4
Patient #48 — $R^2$ = 0.635
3.1
2.4
1.6
0.8
-0.1
1.4
2.9
4.5

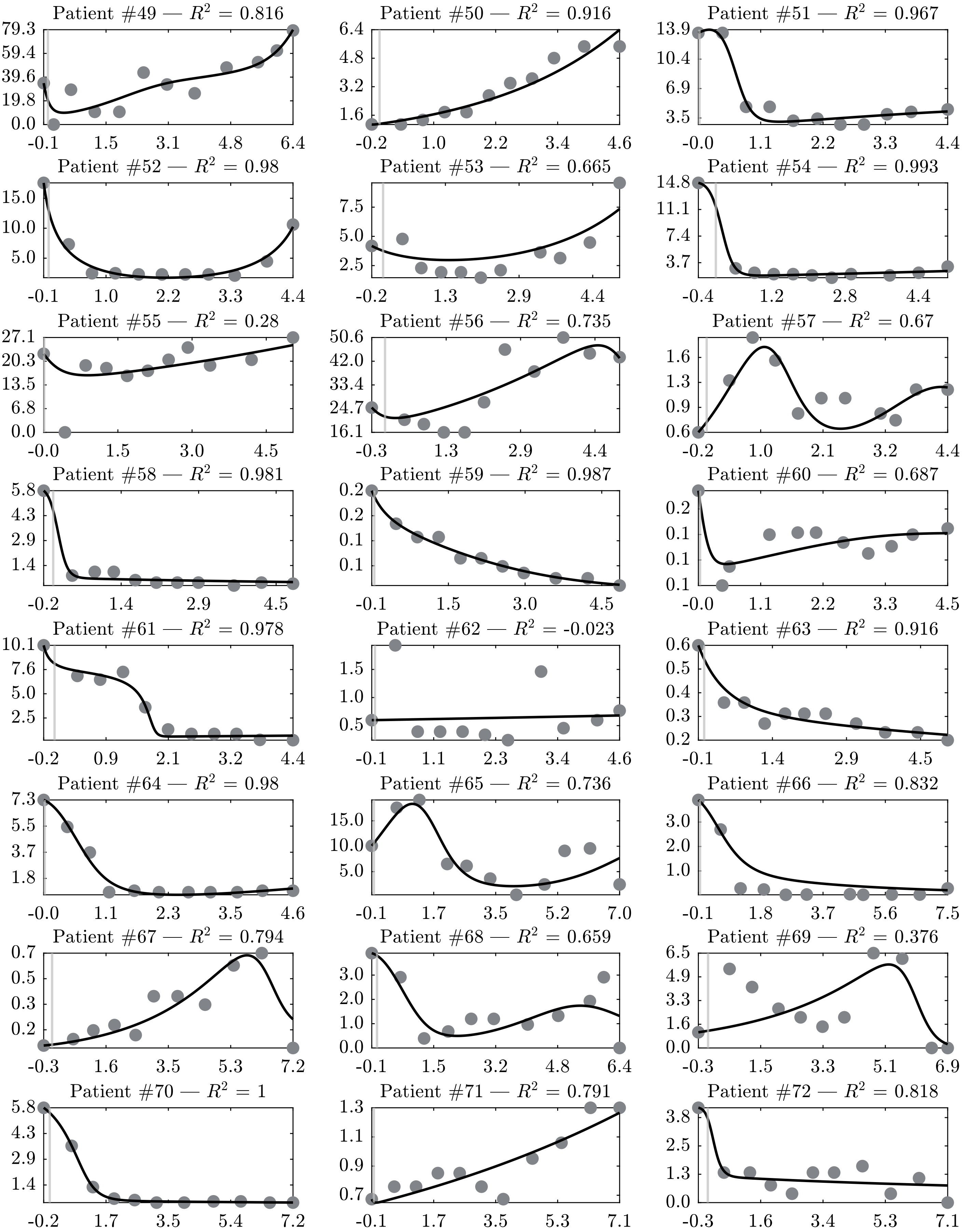
Patient #49 — $R^2$ = 0.816
79.3 59.4 39.6 19.8 0.0
-0.1 1.5 3.1 4.8 6.4
Patient #50 — $R^2$ = 0.916
6.4 4.8 3.2 1.6
-0.2 1.0 2.2 3.4 4.6
Patient #51 — $R^2$ = 0.967
13.9 10.4 6.9 3.5
-0.0 1.1 2.2 3.3 4.4
Patient #52 — $R^2$ = 0.98
15.0 10.0 5.0
-0.1 1.0 2.2 3.3 4.4
Patient #53 — $R^2$ = 0.665
7.5 5.0 2.5
-0.2 1.3 2.9 4.4
Patient #54 — $R^2$ = 0.993
14.8 11.1 7.4 3.7
-0.4 1.2 2.8 4.4
Patient #55 — $R^2$ = 0.28
27.1 20.3 13.5 6.8 0.0
-0.0 1.5 3.0 4.5
Patient #56 — $R^2$ = 0.735
50.6 42.0 33.4 24.7 16.1
-0.3 1.3 2.9 4.4
Patient #57 — $R^2$ = 0.67
1.6 1.3 0.9 0.6
-0.2 1.0 2.1 3.2 4.4
Patient #58 — $R^2$ = 0.981
5.8 4.3 2.9 1.4
-0.2 1.4 2.9 4.5
Patient #59 — $R^2$ = 0.987
0.2 0.2 0.1 0.1
-0.1 1.5 3.0 4.5
Patient #60 — $R^2$ = 0.687
0.2 0.1 0.1 0.1
-0.0 1.1 2.2 3.3 4.5
Patient #61 — $R^2$ = 0.978
10.1 7.6 5.0 2.5
-0.2 0.9 2.1 3.2 4.4
Patient #62 — $R^2$ = -0.023
1.5 1.0 0.5
-0.1 1.1 2.3 3.4 4.6
Patient #63 — $R^2$ = 0.916
0.6 0.5 0.4 0.3 0.2
-0.1 1.4 2.9 4.5
Patient #64 — $R^2$ = 0.98
7.3 5.5 3.7 1.8
-0.0 1.1 2.3 3.5 4.6
Patient #65 — $R^2$ = 0.736
15.0 10.0 5.0
-0.1 1.7 3.5 5.2 7.0
Patient #66 — $R^2$ = 0.832
3.0 2.0 1.0
-0.1 1.8 3.7 5.6 7.5
Patient #67 — $R^2$ = 0.794
0.7 0.5 0.3 0.2
-0.3 1.6 3.5 5.3 7.2
Patient #68 — $R^2$ = 0.659
3.0 2.0 1.0 0.0
-0.1 1.5 3.2 4.8 6.4
Patient #69 — $R^2$ = 0.376
6.5 4.9 3.3 1.6 0.0
-0.3 1.5 3.3 5.1 6.9
Patient #70 — $R^2$ = 1
5.8 4.3 2.9 1.4
-0.2 1.7 3.5 5.4 7.2
Patient #71 — $R^2$ = 0.791
1.3 1.1 0.9 0.7
-0.1 1.7 3.5 5.3 7.1
Patient #72 — $R^2$ = 0.818
3.8 2.5 1.3 0.0
-0.3 1.6 3.4 5.3 7.1

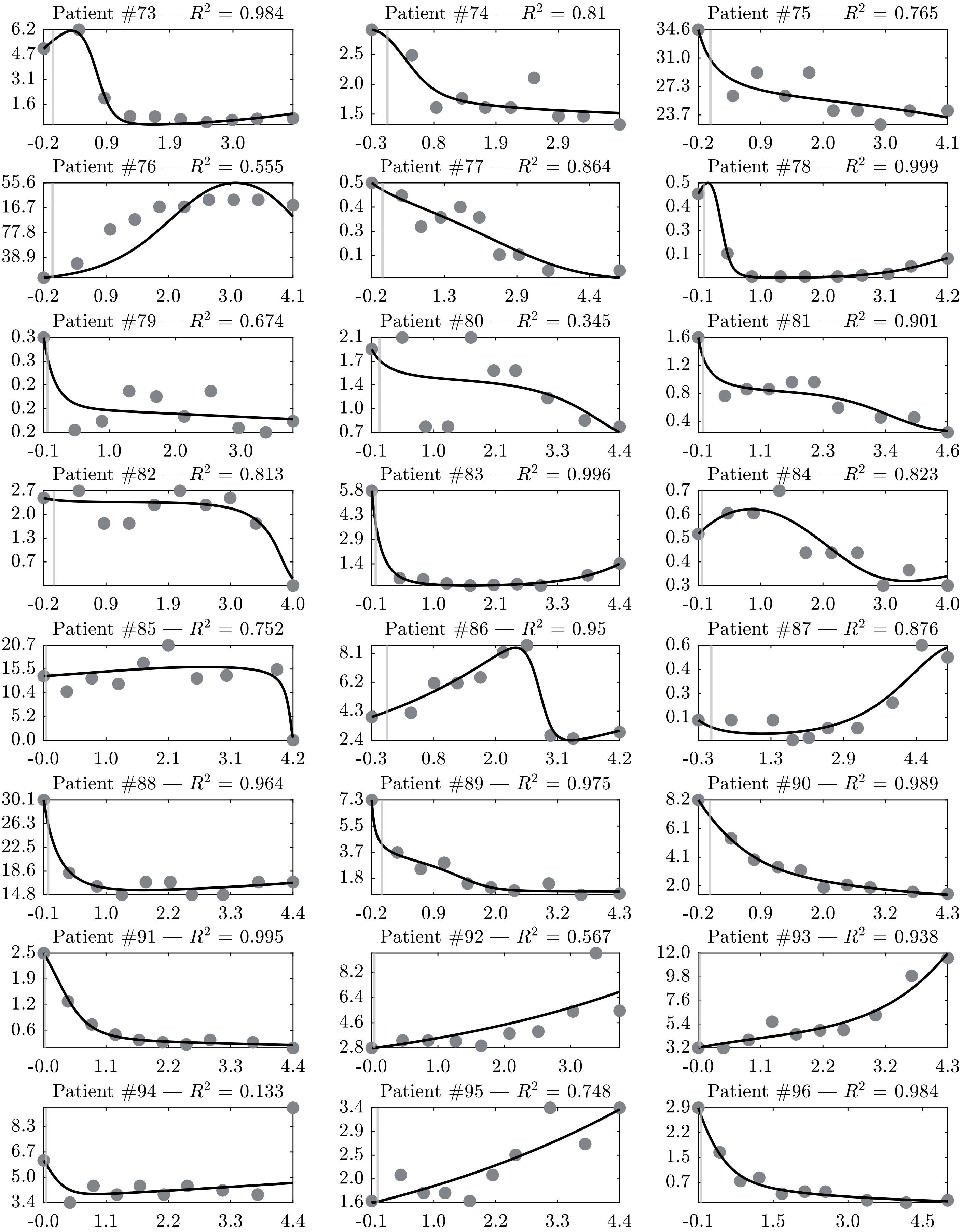

Patient #73 — $R^2$ = 0.984
6.2
4.7
3.1
1.6
-0.2
0.9
1.9
3.0
Patient #74 — $R^2$ = 0.81
2.5
2.0
1.5
-0.3
0.8
1.9
2.9
Patient #75 — $R^2$ = 0.765
34.6
31.0
27.3
23.7
-0.2
0.9
2.0
3.0
4.1
Patient #76 — $R^2$ = 0.555
155.6
116.7
77.8
38.9
-0.2
0.9
2.0
3.0
4.1
Patient #77 — $R^2$ = 0.864
0.5
0.4
0.3
0.1
-0.2
1.3
2.9
4.4
Patient #78 — $R^2$ = 0.999
0.5
0.4
0.3
0.1
-0.1
1.0
2.0
3.1
4.2
Patient #79 — $R^2$ = 0.674
0.3
0.3
0.2
0.2
0.2
-0.1
1.0
2.0
3.0
Patient #80 — $R^2$ = 0.345
2.1
1.7
1.4
1.0
0.7
-0.1
1.0
2.1
3.3
4.4
Patient #81 — $R^2$ = 0.901
1.6
1.2
0.8
0.4
-0.1
1.1
2.3
3.4
4.6
Patient #82 — $R^2$ = 0.813
2.7
2.0
1.3
0.7
-0.2
0.9
1.9
3.0
4.0
Patient #83 — $R^2$ = 0.996
5.8
4.3
2.9
1.4
-0.1
1.0
2.1
3.3
4.4
Patient #84 — $R^2$ = 0.823
0.7
0.6
0.5
0.4
0.3
-0.1
1.0
2.0
3.0
4.0
Patient #85 — $R^2$ = 0.752
20.7
15.5
10.4
5.2
0.0
-0.0
1.0
2.1
3.1
4.2
Patient #86 — $R^2$ = 0.95
8.1
6.2
4.3
2.4
-0.3
0.8
2.0
3.1
4.2
Patient #87 — $R^2$ = 0.876
0.6
0.4
0.3
0.1
-0.3
1.3
2.9
4.4
Patient #88 — $R^2$ = 0.964
30.1
26.3
22.5
18.6
14.8
-0.1
1.0
2.2
3.3
4.4
Patient #89 — $R^2$ = 0.975
7.3
5.5
3.7
1.8
-0.2
0.9
2.0
3.2
4.3
Patient #90 — $R^2$ = 0.989
8.2
6.1
4.1
2.0
-0.2
0.9
2.0
3.2
4.3
Patient #91 — $R^2$ = 0.995
2.5
1.9
1.2
0.6
-0.0
1.1
2.2
3.3
4.4
Patient #92 — $R^2$ = 0.567
8.2
6.4
4.6
2.8
-0.0
1.0
2.0
3.0
Patient #93 — $R^2$ = 0.938
12.0
9.8
7.6
5.4
3.2
-0.0
1.1
2.2
3.2
4.3
Patient #94 — $R^2$ = 0.133
8.3
6.7
5.0
3.4
-0.0
1.1
2.2
3.3
4.4
Patient #95 — $R^2$ = 0.748
3.4
2.9
2.5
2.0
1.6
-0.1
1.0
2.2
3.3
4.4
Patient #96 — $R^2$ = 0.984
2.9
2.2
1.5
0.7
-0.1
1.5
3.0
4.5

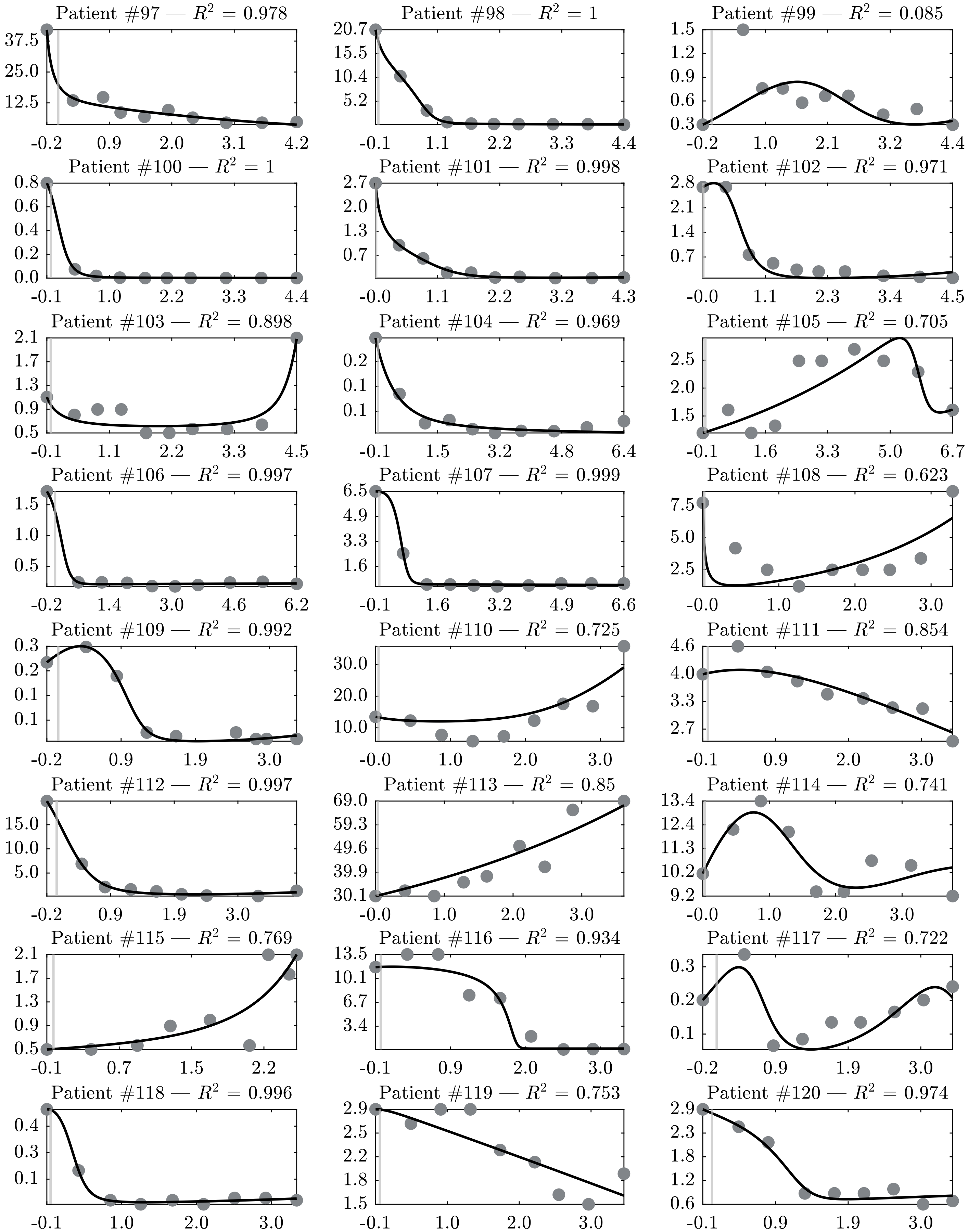
Patient #97 — $R^2$ = 0.978
37.5 25.0 12.5
-0.2 0.9 2.0 3.1 4.2
Patient #98 — $R^2$ = 1
20.7 15.5 10.4 5.2
-0.1 1.1 2.2 3.3 4.4
Patient #99 — $R^2$ = 0.085
1.5 1.2 0.9 0.6 0.3
-0.2 1.0 2.1 3.2 4.4
Patient #100 — $R^2$ = 1
0.8 0.6 0.4 0.2 0.0
-0.1 1.0 2.2 3.3 4.4
Patient #101 — $R^2$ = 0.998
2.7 2.0 1.3 0.7
-0.0 1.1 2.2 3.2 4.3
Patient #102 — $R^2$ = 0.971
2.8 2.1 1.4 0.7
-0.0 1.1 2.3 3.4 4.5
Patient #103 — $R^2$ = 0.898
2.1 1.7 1.3 0.9 0.5
-0.1 1.1 2.2 3.3 4.5
Patient #104 — $R^2$ = 0.969
0.2 0.1 0.1
-0.1 1.5 3.2 4.8 6.4
Patient #105 — $R^2$ = 0.705
2.5 2.0 1.5
-0.1 1.6 3.3 5.0 6.7
Patient #106 — $R^2$ = 0.997
1.5 1.0 0.5
-0.2 1.4 3.0 4.6 6.2
Patient #107 — $R^2$ = 0.999
6.5 4.9 3.3 1.6
-0.1 1.6 3.2 4.9 6.6
Patient #108 — $R^2$ = 0.623
7.5 5.0 2.5
-0.0 1.0 2.0 3.0
Patient #109 — $R^2$ = 0.992
0.3 0.2 0.1 0.1
-0.2 0.9 1.9 3.0
Patient #110 — $R^2$ = 0.725
30.0 20.0 10.0
-0.0 1.0 2.0 3.0
Patient #111 — $R^2$ = 0.854
4.6 4.0 3.3 2.7
-0.1 0.9 2.0 3.0
Patient #112 — $R^2$ = 0.997
15.0 10.0 5.0
-0.2 0.9 1.9 3.0
Patient #113 — $R^2$ = 0.85
69.0 59.3 49.6 39.9 30.1
-0.0 1.0 2.0 3.0
Patient #114 — $R^2$ = 0.741
13.4 12.4 11.3 10.2 9.2
-0.0 1.0 2.0 3.0
Patient #115 — $R^2$ = 0.769
2.1 1.7 1.3 0.9 0.5
-0.1 0.7 1.5 2.2
Patient #116 — $R^2$ = 0.934
13.5 10.1 6.7 3.4
-0.1 0.9 2.0 3.0
Patient #117 — $R^2$ = 0.722
0.3 0.2 0.1
-0.2 0.9 1.9 3.0
Patient #118 — $R^2$ = 0.996
0.4 0.3 0.1
-0.1 1.0 2.0 3.0
Patient #119 — $R^2$ = 0.753
2.9 2.5 2.2 1.8 1.5
-0.1 1.0 2.0 3.0
Patient #120 — $R^2$ = 0.974
2.9 2.3 1.8 1.2 0.6
-0.1 0.9 1.9 3.0

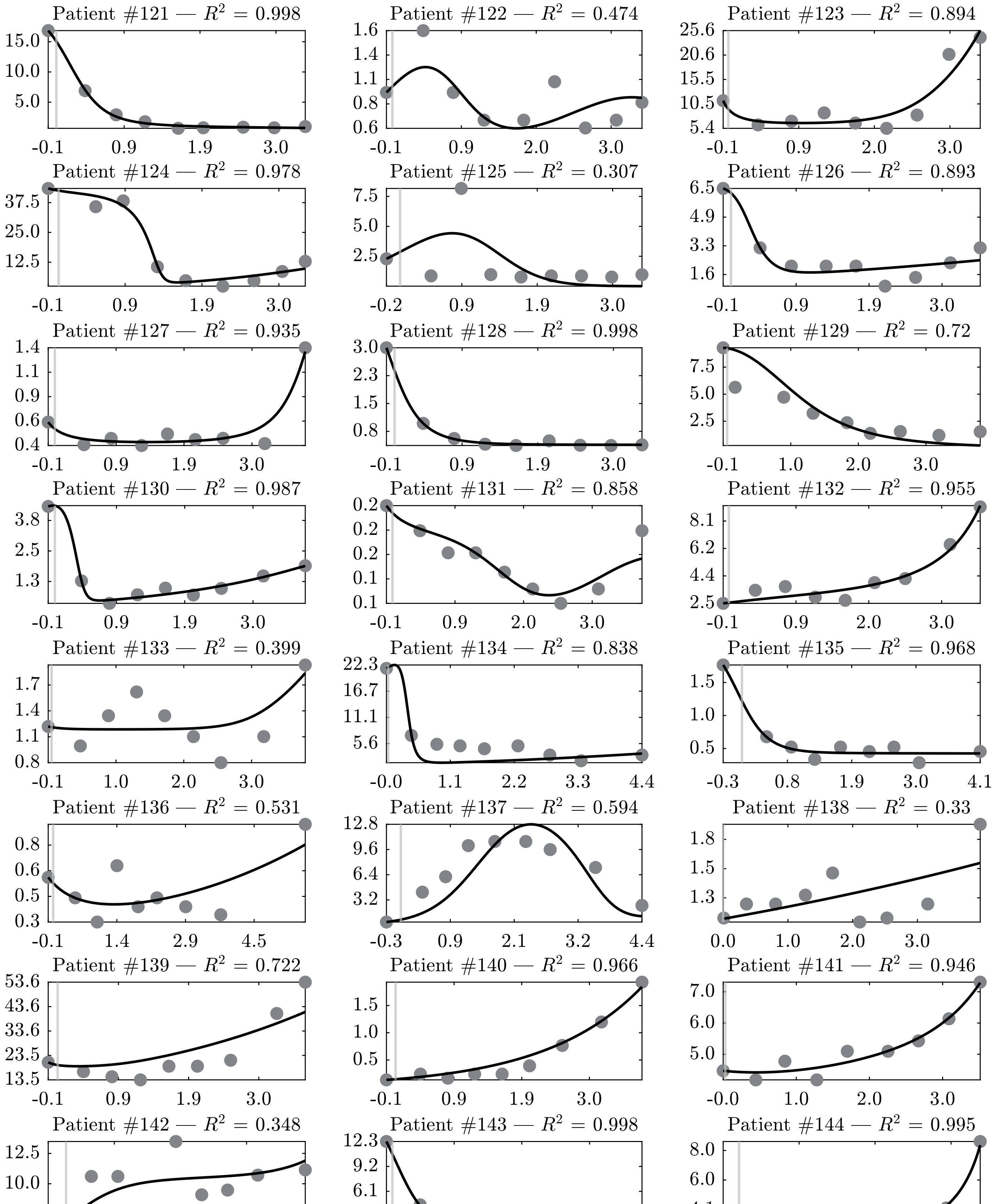

Patient #121 — $R^2$ = 0.998
15.0 10.0 5.0
-0.1 0.9 1.9 3.0
Patient #122 — $R^2$ = 0.474
1.6 1.4 1.1 0.8 0.6
-0.1 0.9 2.0 3.0
Patient #123 — $R^2$ = 0.894
25.6 20.6 15.5 10.5 5.4
-0.1 0.9 2.0 3.0
Patient #124 — $R^2$ = 0.978
37.5 25.0 12.5
-0.1 0.9 1.9 3.0
Patient #125 — $R^2$ = 0.307
7.5 5.0 2.5
-0.2 0.9 1.9 3.0
Patient #126 — $R^2$ = 0.893
6.5 4.9 3.3 1.6
-0.1 0.9 1.9 3.0
Patient #127 — $R^2$ = 0.935
1.4 1.1 0.9 0.6 0.4
-0.1 0.9 1.9 3.0
Patient #128 — $R^2$ = 0.998
3.0 2.3 1.5 0.8
-0.1 0.9 1.9 3.0
Patient #129 — $R^2$ = 0.72
7.5 5.0 2.5
-0.1 1.0 2.0 3.0
Patient #130 — $R^2$ = 0.987
3.8 2.5 1.3
-0.1 0.9 1.9 3.0
Patient #131 — $R^2$ = 0.858
0.2 0.2 0.2 0.1 0.1
-0.1 0.9 2.0 3.0
Patient #132 — $R^2$ = 0.955
8.1 6.2 4.4 2.5
-0.1 0.9 2.0 3.0
Patient #133 — $R^2$ = 0.399
1.7 1.4 1.1 0.8
-0.1 1.0 2.0 3.0
Patient #134 — $R^2$ = 0.838
22.3 16.7 11.1 5.6
-0.0 1.1 2.2 3.3 4.4
Patient #135 — $R^2$ = 0.968
1.5 1.0 0.5
-0.3 0.8 1.9 3.0 4.1
Patient #136 — $R^2$ = 0.531
0.8 0.6 0.5 0.3
-0.1 1.4 2.9 4.5
Patient #137 — $R^2$ = 0.594
12.8 9.6 6.4 3.2
-0.3 0.9 2.1 3.2 4.4
Patient #138 — $R^2$ = 0.33
1.8 1.5 1.3
0.0 1.0 2.0 3.0
Patient #139 — $R^2$ = 0.722
53.6 43.6 33.6 23.5 13.5
-0.1 0.9 1.9 3.0
Patient #140 — $R^2$ = 0.966
1.5 1.0 0.5
-0.1 0.9 1.9 3.0
Patient #141 — $R^2$ = 0.946
7.0 6.0 5.0
-0.0 1.0 2.0 3.0
Patient #142 — $R^2$ = 0.348
12.5 10.0 7.5
-0.3 0.8 1.9 2.9
Patient #143 — $R^2$ = 0.998
12.3 9.2 6.1 3.1
-0.1 0.9 2.0 3.0
Patient #144 — $R^2$ = 0.995
8.0 6.0 4.1 2.1
-0.2 0.8 1.9 2.9

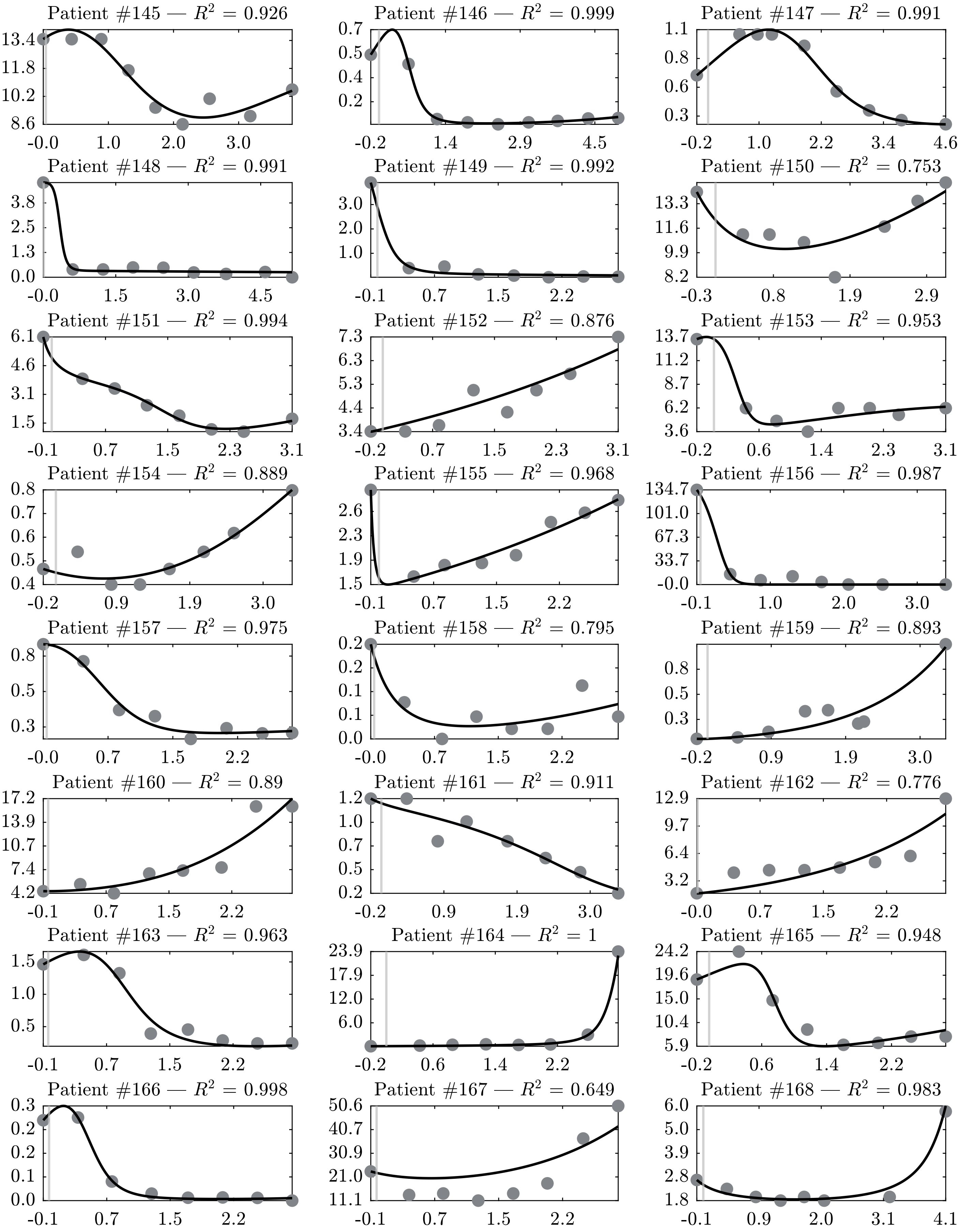
Patient #145 — $R^2$ = 0.926
13.4 11.8 10.2 8.6
-0.0 1.0 2.0 3.0
Patient #146 — $R^2$ = 0.999
0.7 0.5 0.4 0.2
-0.2 1.4 2.9 4.5
Patient #147 — $R^2$ = 0.991
1.1 0.8 0.6 0.3
-0.2 1.0 2.2 3.4 4.6
Patient #148 — $R^2$ = 0.991
3.8 2.5 1.3 0.0
-0.0 1.5 3.0 4.5
Patient #149 — $R^2$ = 0.992
3.0 2.0 1.0
-0.1 0.7 1.5 2.2
Patient #150 — $R^2$ = 0.753
13.3 11.6 9.9 8.2
-0.3 0.8 1.9 2.9
Patient #151 — $R^2$ = 0.994
6.1 4.6 3.1 1.5
-0.1 0.7 1.5 2.3 3.1
Patient #152 — $R^2$ = 0.876
7.3 6.3 5.3 4.4 3.4
-0.2 0.7 1.5 2.3 3.1
Patient #153 — $R^2$ = 0.953
13.7 11.2 8.7 6.2 3.6
-0.2 0.6 1.4 2.3 3.1
Patient #154 — $R^2$ = 0.889
0.8 0.7 0.6 0.5 0.4
-0.2 0.9 1.9 3.0
Patient #155 — $R^2$ = 0.968
2.6 2.3 1.9 1.5
-0.1 0.7 1.5 2.2
Patient #156 — $R^2$ = 0.987
134.7 101.0 67.3 33.7 -0.0
-0.1 1.0 2.0 3.0
Patient #157 — $R^2$ = 0.975
0.8 0.5 0.3
-0.0 0.7 1.5 2.2
Patient #158 — $R^2$ = 0.795
0.2 0.2 0.1 0.1 0.0
0.0 0.7 1.5 2.2
Patient #159 — $R^2$ = 0.893
0.8 0.5 0.3
-0.2 0.9 1.9 3.0
Patient #160 — $R^2$ = 0.89
17.2 13.9 10.7 7.4 4.2
-0.1 0.7 1.5 2.2
Patient #161 — $R^2$ = 0.911
1.2 1.0 0.7 0.5 0.2
-0.2 0.9 1.9 3.0
Patient #162 — $R^2$ = 0.776
12.9 9.7 6.4 3.2
-0.0 0.7 1.5 2.2
Patient #163 — $R^2$ = 0.963
1.5 1.0 0.5
-0.1 0.7 1.5 2.2
Patient #164 — $R^2$ = 1
23.9 17.9 12.0 6.0
-0.2 0.6 1.4 2.2
Patient #165 — $R^2$ = 0.948
24.2 19.6 15.0 10.4 5.9
-0.2 0.6 1.4 2.2
Patient #166 — $R^2$ = 0.998
0.3 0.2 0.2 0.1 0.0
-0.1 0.7 1.5 2.2
Patient #167 — $R^2$ = 0.649
50.6 40.7 30.9 21.0 11.1
-0.1 0.7 1.5 2.2
Patient #168 — $R^2$ = 0.983
6.0 5.0 3.9 2.8 1.8
-0.1 0.9 2.0 3.1 4.1

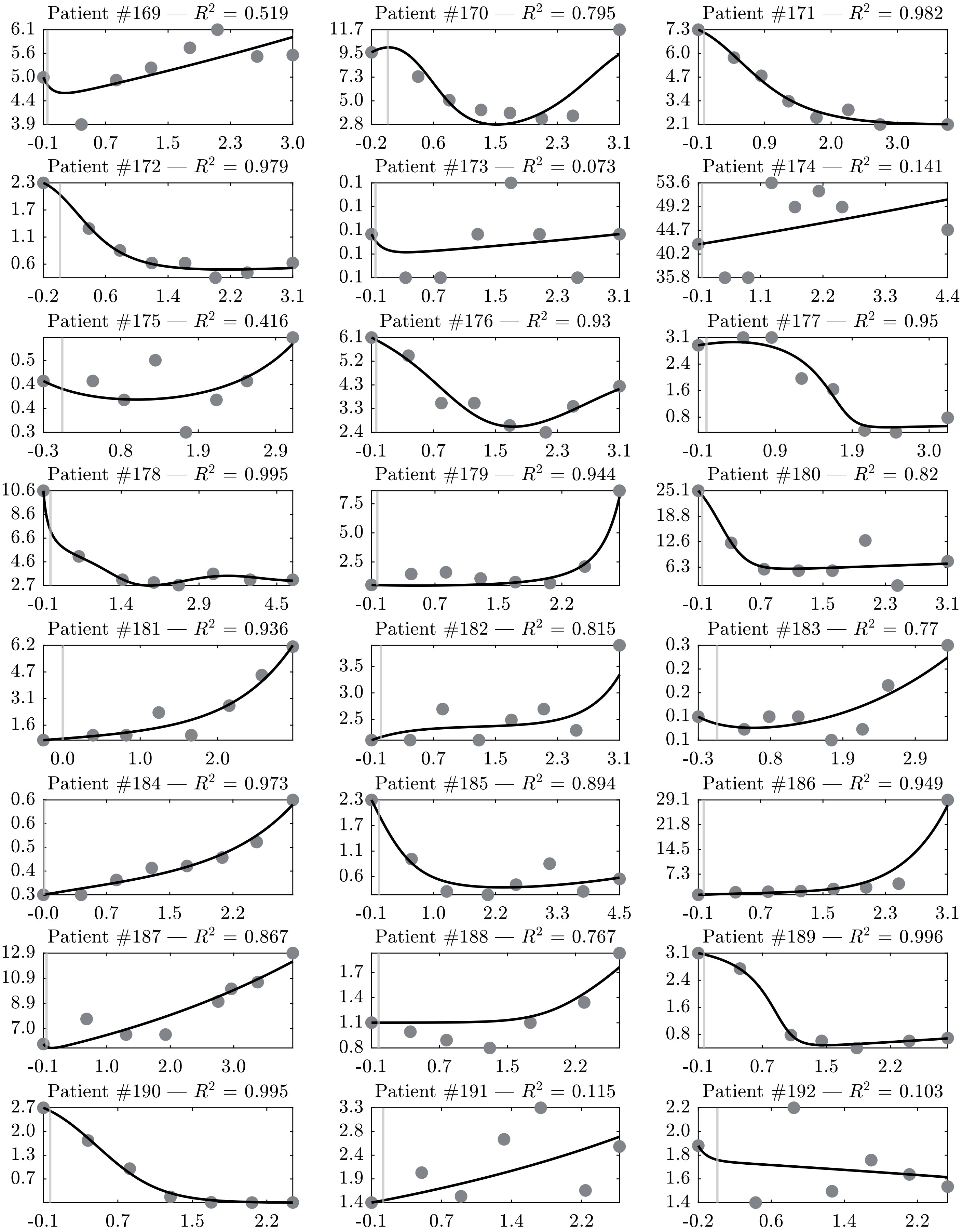
Patient #169 — $R^2$ = 0.519
6.1 5.6 5.0 4.4 3.9
-0.1 0.7 1.5 2.3 3.0
Patient #170 — $R^2$ = 0.795
11.7 9.5 7.3 5.0 2.8
-0.2 0.6 1.5 2.3 3.1
Patient #171 — $R^2$ = 0.982
7.3 6.0 4.7 3.4 2.1
-0.1 0.9 2.0 3.0
Patient #172 — $R^2$ = 0.979
2.3 1.7 1.1 0.6
-0.2 0.6 1.4 2.2 3.1
Patient #173 — $R^2$ = 0.073
0.1 0.1 0.1 0.1 0.1
-0.1 0.7 1.5 2.3 3.1
Patient #174 — $R^2$ = 0.141
53.6 49.2 44.7 40.2 35.8
-0.1 1.1 2.2 3.3 4.4
Patient #175 — $R^2$ = 0.416
0.5 0.4 0.4 0.3
-0.3 0.8 1.9 2.9
Patient #176 — $R^2$ = 0.93
6.1 5.2 4.3 3.3 2.4
-0.1 0.7 1.5 2.3 3.1
Patient #177 — $R^2$ = 0.95
3.1 2.4 1.6 0.8
-0.1 0.9 1.9 3.0
Patient #178 — $R^2$ = 0.995
10.6 8.6 6.6 4.6 2.7
-0.1 1.4 2.9 4.5
Patient #179 — $R^2$ = 0.944
7.5 5.0 2.5
-0.1 0.7 1.5 2.2
Patient #180 — $R^2$ = 0.82
25.1 18.8 12.6 6.3
-0.1 0.7 1.5 2.3 3.1
Patient #181 — $R^2$ = 0.936
6.2 4.7 3.1 1.6
0.0 1.0 2.0
Patient #182 — $R^2$ = 0.815
3.5 3.0 2.5
-0.1 0.7 1.5 2.3 3.1
Patient #183 — $R^2$ = 0.77
0.3 0.2 0.2 0.1 0.1
-0.3 0.8 1.9 2.9
Patient #184 — $R^2$ = 0.973
0.6 0.6 0.5 0.4 0.3
-0.0 0.7 1.5 2.2
Patient #185 — $R^2$ = 0.894
2.3 1.7 1.1 0.6
-0.1 1.0 2.2 3.3 4.5
Patient #186 — $R^2$ = 0.949
29.1 21.8 14.5 7.3
-0.1 0.7 1.5 2.3 3.1
Patient #187 — $R^2$ = 0.867
12.9 10.9 8.9 7.0
-0.1 1.0 2.0 3.0
Patient #188 — $R^2$ = 0.767
1.7 1.4 1.1 0.8
-0.1 0.7 1.5 2.2
Patient #189 — $R^2$ = 0.996
3.1 2.4 1.6 0.8
-0.1 0.7 1.5 2.2
Patient #190 — $R^2$ = 0.995
2.7 2.0 1.3 0.7
-0.1 0.7 1.5 2.2
Patient #191 — $R^2$ = 0.115
3.3 2.8 2.4 1.9 1.4
-0.1 0.7 1.4 2.2
Patient #192 — $R^2$ = 0.103
2.2 2.0 1.8 1.6 1.4
-0.2 0.6 1.4 2.2

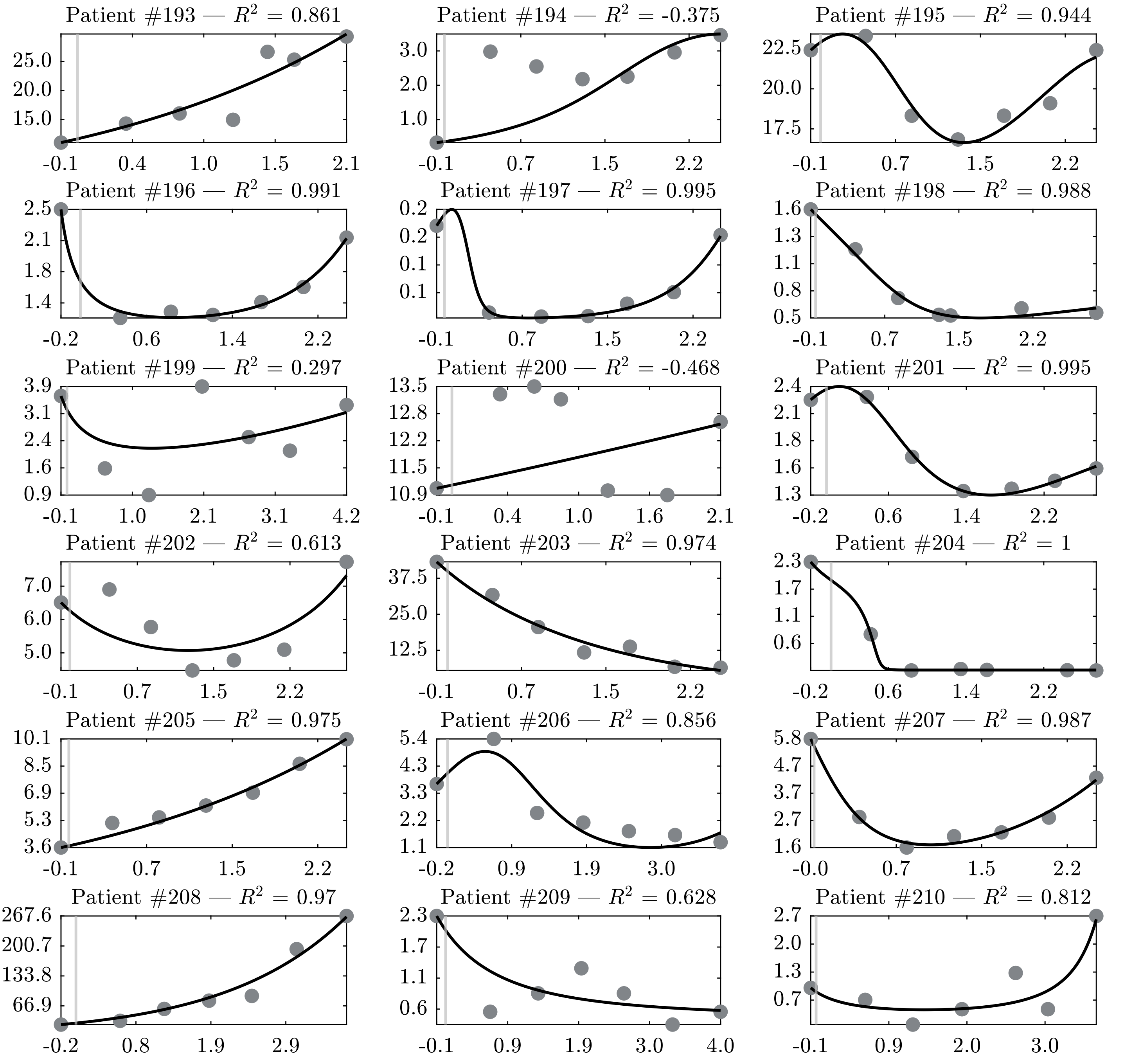
Patient #193 — $R^2$ = 0.861
25.0
20.0
15.0
-0.1
0.4
1.0
1.5
2.1
Patient #194 — $R^2$ = -0.375
3.0
2.0
1.0
-0.1
0.7
1.5
2.2
Patient #195 — $R^2$ = 0.944
22.5
20.0
17.5
-0.1
0.7
1.5
2.2
Patient #196 — $R^2$ = 0.991
2.5
2.1
1.8
1.4
-0.2
0.6
1.4
2.2
Patient #197 — $R^2$ = 0.995
0.2
0.2
0.1
0.1
-0.1
0.7
1.5
2.2
Patient #198 — $R^2$ = 0.988
1.6
1.3
1.1
0.8
0.5
-0.1
0.7
1.5
2.2
Patient #199 — $R^2$ = 0.297
3.9
3.1
2.4
1.6
0.9
-0.1
1.0
2.1
3.1
4.2
Patient #200 — $R^2$ = -0.468
13.5
12.8
12.2
11.5
10.9
-0.1
0.4
1.0
1.6
2.1
Patient #201 — $R^2$ = 0.995
2.4
2.1
1.8
1.6
1.3
-0.2
0.6
1.4
2.2
Patient #202 — $R^2$ = 0.613
7.0
6.0
5.0
-0.1
0.7
1.5
2.2
Patient #203 — $R^2$ = 0.974
37.5
25.0
12.5
-0.1
0.7
1.5
2.2
Patient #204 — $R^2$ = 1
2.3
1.7
1.1
0.6
-0.2
0.6
1.4
2.2
Patient #205 — $R^2$ = 0.975
10.1
8.5
6.9
5.3
3.6
-0.1
0.7
1.5
2.2
Patient #206 — $R^2$ = 0.856
5.4
4.3
3.3
2.2
1.1
-0.2
0.9
1.9
3.0
Patient #207 — $R^2$ = 0.987
5.8
4.7
3.7
2.7
1.6
-0.0
0.7
1.5
2.2
Patient #208 — $R^2$ = 0.97
267.6
200.7
133.8
66.9
-0.2
0.8
1.9
2.9
Patient #209 — $R^2$ = 0.628
2.3
1.7
1.1
0.6
-0.1
0.9
1.9
3.0
4.0
Patient #210 — $R^2$ = 0.812
2.7
2.0
1.3
0.7
-0.1
0.9
2.0
3.0

Model extrapolation results of all 210 patients. The solid-line curves and the points represent model results and measured data, respectively. The last two data points are not used for parameters estimation. Ordinates: normalized number of tumor cells. Abscissas: normalized treatment time, negative values indicate time before the start of treatment. The model is capable of forecasting tumor dynamics qualitatively and sometimes quantitatively.

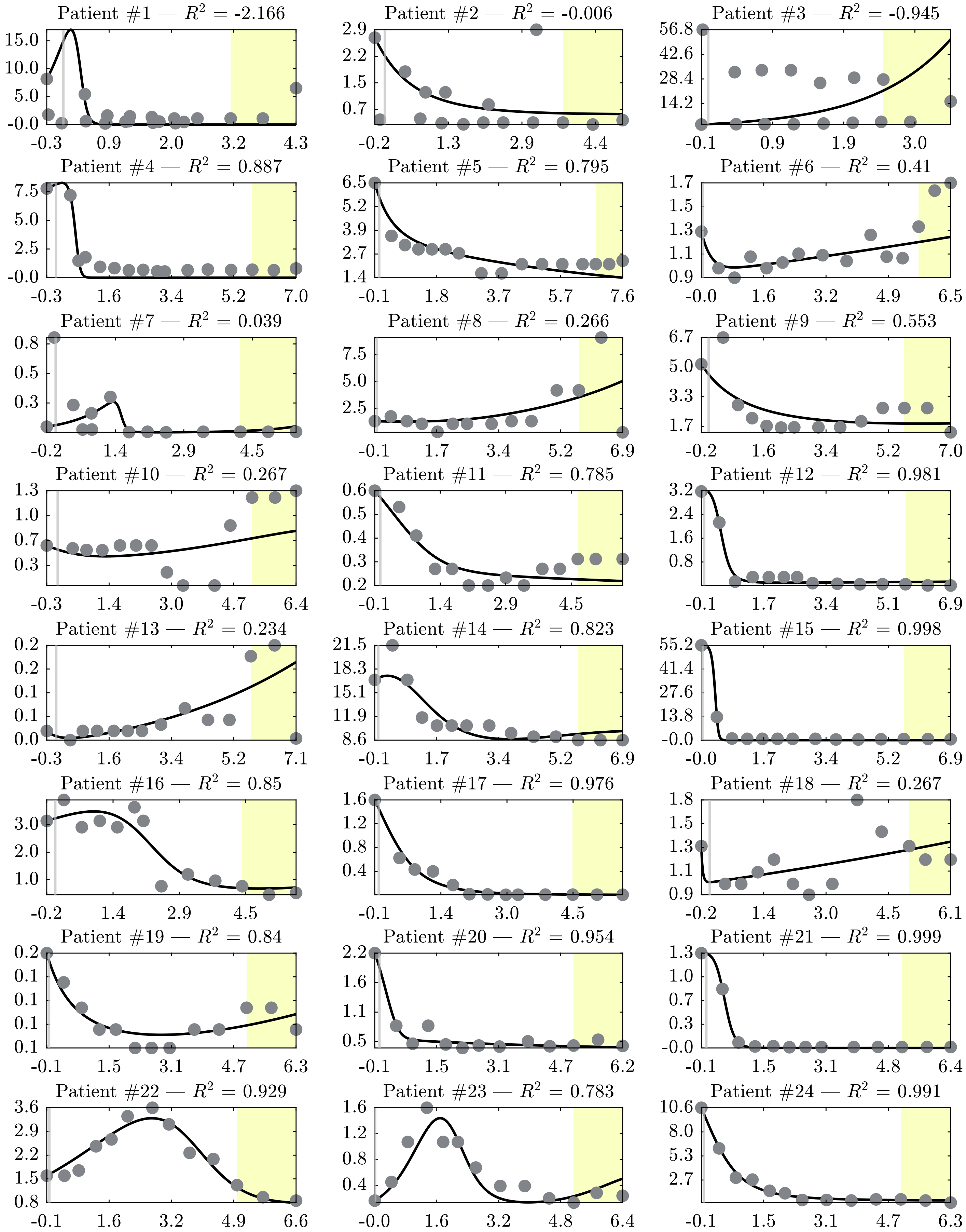

Patient #1 — $R^2$ = -2.166
15.0 10.0 5.0 -0.0
-0.3 0.9 2.0 3.2 4.3
Patient #2 — $R^2$ = -0.006
2.9 2.2 1.5 0.7
-0.2 1.3 2.9 4.4
Patient #3 — $R^2$ = -0.945
56.8 42.6 28.4 14.2
-0.1 0.9 1.9 3.0
Patient #4 — $R^2$ = 0.887
7.5 5.0 2.5 -0.0
-0.3 1.6 3.4 5.2 7.0
Patient #5 — $R^2$ = 0.795
6.5 5.2 3.9 2.7 1.4
-0.1 1.8 3.7 5.7 7.6
Patient #6 — $R^2$ = 0.41
1.7 1.5 1.3 1.1 0.9
-0.0 1.6 3.2 4.9 6.5
Patient #7 — $R^2$ = 0.039
0.8 0.5 0.3
-0.2 1.4 2.9 4.5
Patient #8 — $R^2$ = 0.266
7.5 5.0 2.5
-0.1 1.7 3.4 5.2 6.9
Patient #9 — $R^2$ = 0.553
6.7 5.0 3.3 1.7
-0.2 1.6 3.4 5.2 7.0
Patient #10 — $R^2$ = 0.267
1.3 1.0 0.7 0.3
-0.3 1.4 3.0 4.7 6.4
Patient #11 — $R^2$ = 0.785
0.6 0.5 0.4 0.3 0.2
-0.1 1.4 2.9 4.5
Patient #12 — $R^2$ = 0.981
3.2 2.4 1.6 0.8
-0.1 1.7 3.4 5.1 6.9
Patient #13 — $R^2$ = 0.234
0.2 0.2 0.1 0.1 0.0
-0.3 1.6 3.4 5.2 7.1
Patient #14 — $R^2$ = 0.823
21.5 18.3 15.1 11.9 8.6
-0.1 1.7 3.4 5.2 6.9
Patient #15 — $R^2$ = 0.998
55.2 41.4 27.6 13.8 -0.0
-0.0 1.7 3.4 5.2 6.9
Patient #16 — $R^2$ = 0.85
3.0 2.0 1.0
-0.2 1.4 2.9 4.5
Patient #17 — $R^2$ = 0.976
1.6 1.2 0.8 0.4
-0.1 1.4 3.0 4.5
Patient #18 — $R^2$ = 0.267
1.8 1.5 1.3 1.1 0.9
-0.2 1.4 3.0 4.5 6.1
Patient #19 — $R^2$ = 0.84
0.2 0.1 0.1 0.1 0.1
-0.1 1.5 3.1 4.7 6.3
Patient #20 — $R^2$ = 0.954
2.2 1.6 1.1 0.5
-0.1 1.5 3.1 4.7 6.2
Patient #21 — $R^2$ = 0.999
1.3 1.0 0.7 0.3 -0.0
-0.1 1.5 3.1 4.8 6.4
Patient #22 — $R^2$ = 0.929
3.6 2.9 2.2 1.5 0.8
-0.1 1.6 3.2 4.9 6.6
Patient #23 — $R^2$ = 0.783
1.6 1.2 0.8 0.4
-0.0 1.6 3.2 4.8 6.4
Patient #24 — $R^2$ = 0.991
10.6 8.0 5.3 2.7
-0.1 1.5 3.1 4.7 6.3

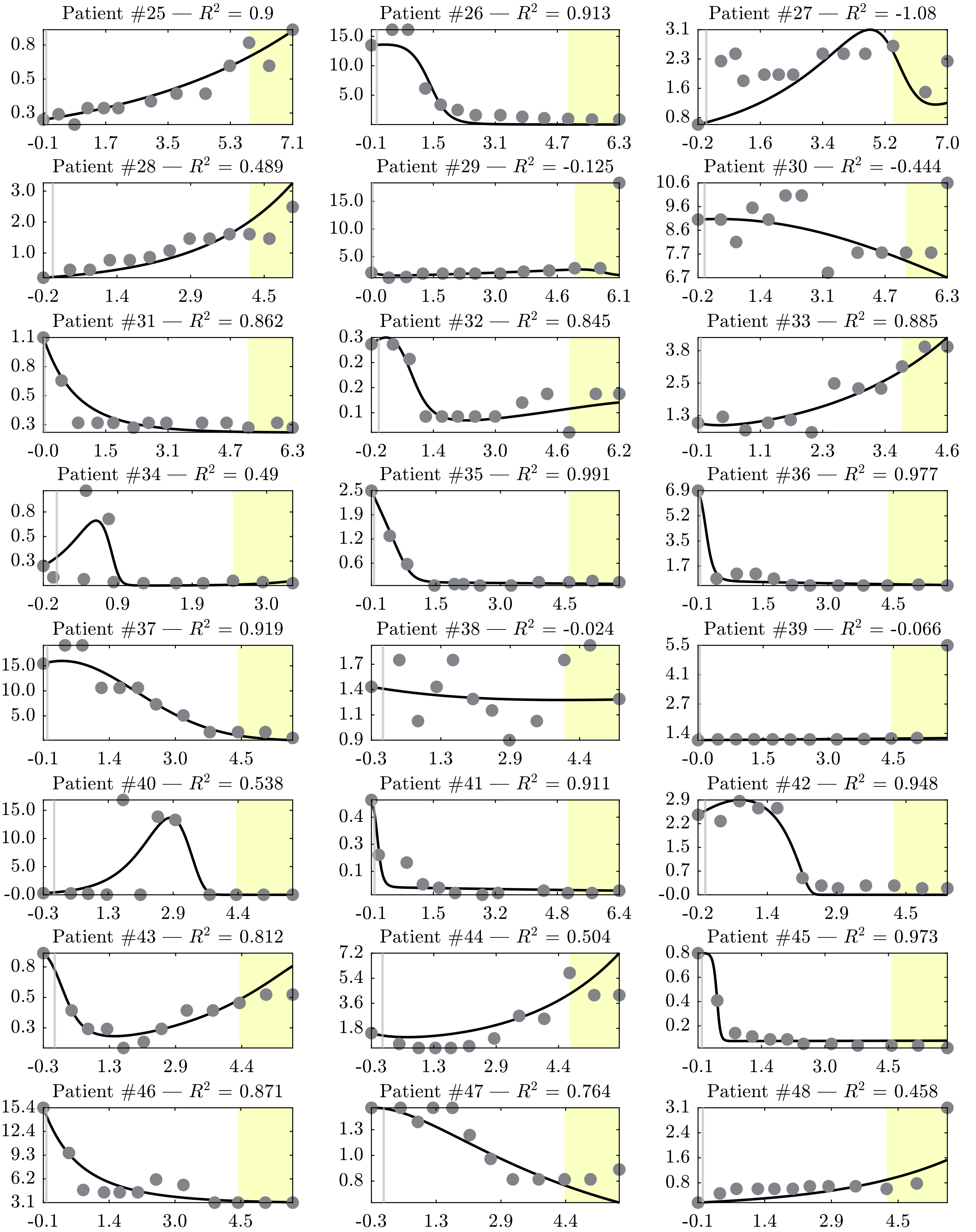
Patient #25 — $R^2$ = 0.9
0.8 0.5 0.3
-0.1 1.7 3.5 5.3 7.1
Patient #26 — $R^2$ = 0.913
15.0 10.0 5.0
-0.1 1.5 3.1 4.7 6.3
Patient #27 — $R^2$ = -1.08
3.1 2.3 1.6 0.8
-0.2 1.6 3.4 5.2 7.0
Patient #28 — $R^2$ = 0.489
3.0 2.0 1.0
-0.2 1.4 2.9 4.5
Patient #29 — $R^2$ = -0.125
15.0 10.0 5.0
-0.0 1.5 3.0 4.6 6.1
Patient #30 — $R^2$ = -0.444
10.6 9.6 8.6 7.7 6.7
-0.2 1.4 3.1 4.7 6.3
Patient #31 — $R^2$ = 0.862
1.1 0.8 0.5 0.3
-0.0 1.5 3.1 4.7 6.3
Patient #32 — $R^2$ = 0.845
0.3 0.2 0.2 0.1
-0.2 1.4 3.0 4.6 6.2
Patient #33 — $R^2$ = 0.885
3.8 2.5 1.3
-0.1 1.1 2.3 3.4 4.6
Patient #34 — $R^2$ = 0.49
0.8 0.5 0.3
-0.2 0.9 1.9 3.0
Patient #35 — $R^2$ = 0.991
2.5 1.9 1.2 0.6
-0.1 1.5 3.0 4.5
Patient #36 — $R^2$ = 0.977
6.9 5.2 3.5 1.7
-0.1 1.5 3.0 4.5
Patient #37 — $R^2$ = 0.919
15.0 10.0 5.0
-0.1 1.4 3.0 4.5
Patient #38 — $R^2$ = -0.024
1.7 1.4 1.1 0.9
-0.3 1.3 2.9 4.4
Patient #39 — $R^2$ = -0.066
5.5 4.1 2.7 1.4
-0.0 1.5 3.0 4.5
Patient #40 — $R^2$ = 0.538
15.0 10.0 5.0 -0.0
-0.3 1.3 2.9 4.4
Patient #41 — $R^2$ = 0.911
0.4 0.3 0.1
-0.1 1.5 3.2 4.8 6.4
Patient #42 — $R^2$ = 0.948
2.9 2.2 1.5 0.7 -0.0
-0.2 1.4 2.9 4.5
Patient #43 — $R^2$ = 0.812
0.8 0.5 0.3
-0.3 1.3 2.9 4.4
Patient #44 — $R^2$ = 0.504
7.2 5.4 3.6 1.8
-0.3 1.3 2.9 4.4
Patient #45 — $R^2$ = 0.973
0.8 0.6 0.4 0.2
-0.1 1.4 3.0 4.5
Patient #46 — $R^2$ = 0.871
15.4 12.4 9.3 6.2 3.1
-0.1 1.4 3.0 4.5
Patient #47 — $R^2$ = 0.764
1.3 1.0 0.8
-0.3 1.3 2.9 4.4
Patient #48 — $R^2$ = 0.458
3.1 2.4 1.6 0.8
-0.1 1.4 2.9 4.5

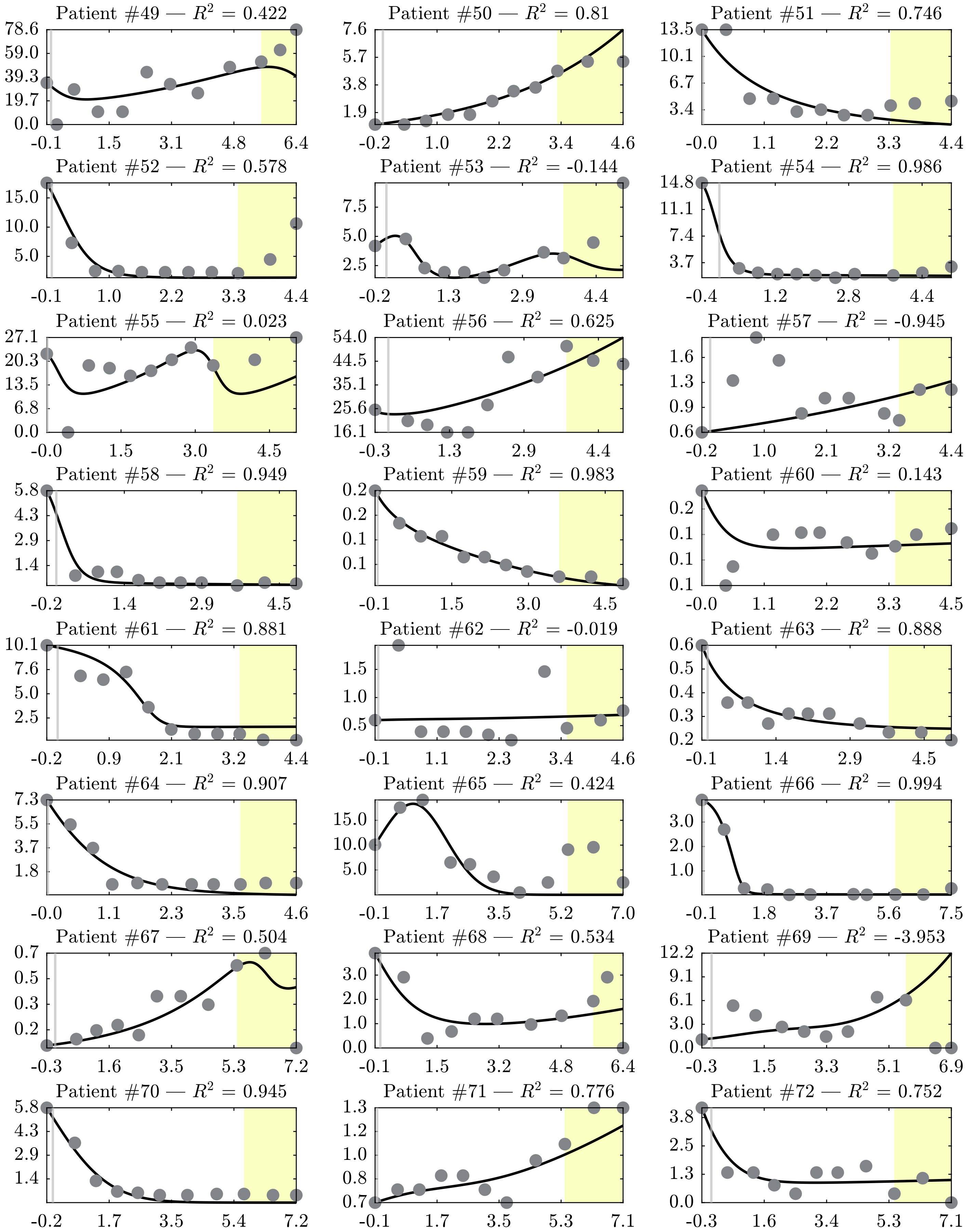

Patient #49 — $R^2$ = 0.422
Patient #50 — $R^2$ = 0.81
Patient #51 — $R^2$ = 0.746
Patient #52 — $R^2$ = 0.578
Patient #53 — $R^2$ = -0.144
Patient #54 — $R^2$ = 0.986
Patient #55 — $R^2$ = 0.023
Patient #56 — $R^2$ = 0.625
Patient #57 — $R^2$ = -0.945
Patient #58 — $R^2$ = 0.949
Patient #59 — $R^2$ = 0.983
Patient #60 — $R^2$ = 0.143
Patient #61 — $R^2$ = 0.881
Patient #62 — $R^2$ = -0.019
Patient #63 — $R^2$ = 0.888
Patient #64 — $R^2$ = 0.907
Patient #65 — $R^2$ = 0.424
Patient #66 — $R^2$ = 0.994
Patient #67 — $R^2$ = 0.504
Patient #68 — $R^2$ = 0.534
Patient #69 — $R^2$ = -3.953
Patient #70 — $R^2$ = 0.945
Patient #71 — $R^2$ = 0.776
Patient #72 — $R^2$ = 0.752

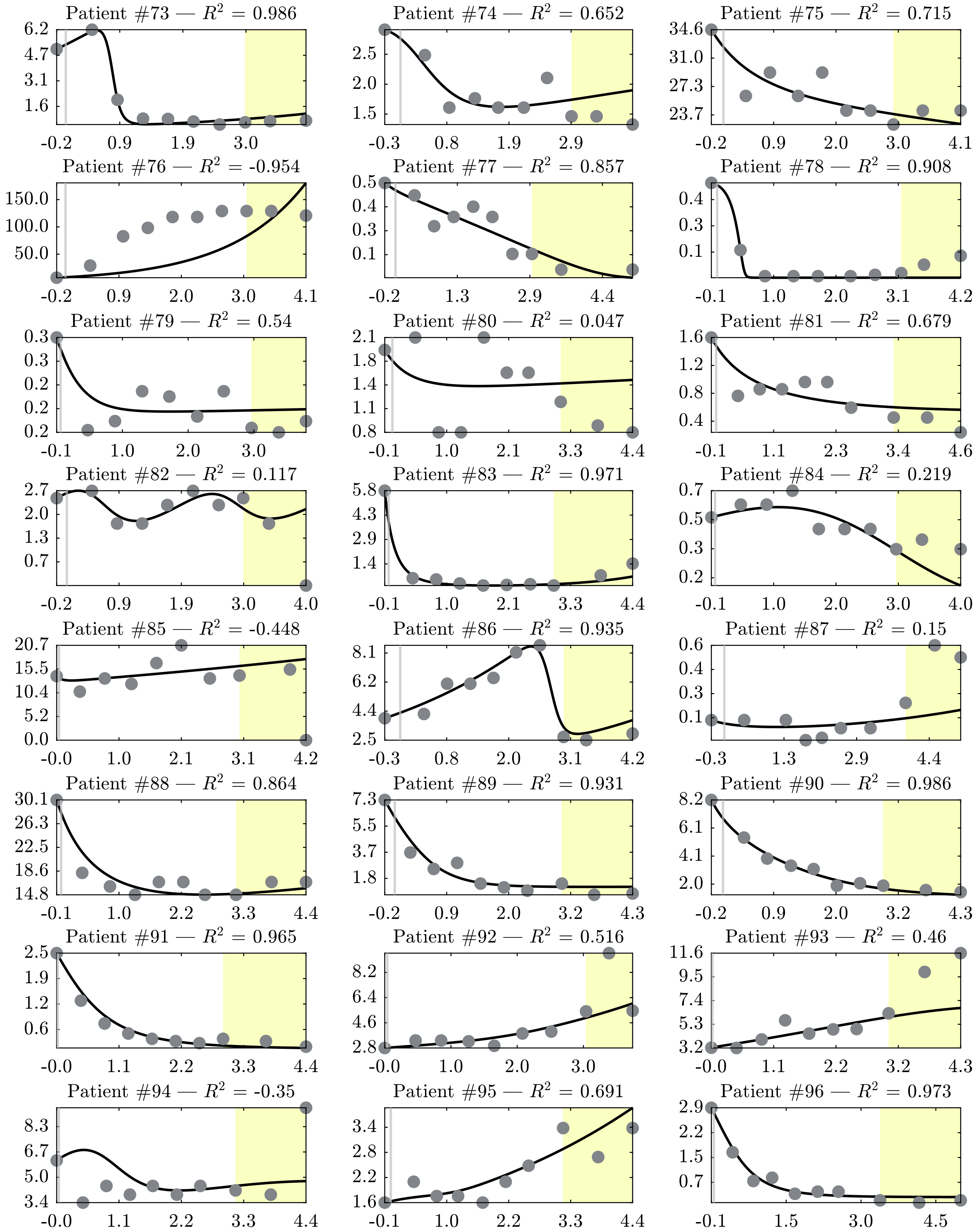

Patient #73 — $R^2$ = 0.986
Patient #74 — $R^2$ = 0.652
Patient #75 — $R^2$ = 0.715
Patient #76 — $R^2$ = -0.954
Patient #77 — $R^2$ = 0.857
Patient #78 — $R^2$ = 0.908
Patient #79 — $R^2$ = 0.54
Patient #80 — $R^2$ = 0.047
Patient #81 — $R^2$ = 0.679
Patient #82 — $R^2$ = 0.117
Patient #83 — $R^2$ = 0.971
Patient #84 — $R^2$ = 0.219
Patient #85 — $R^2$ = -0.448
Patient #86 — $R^2$ = 0.935
Patient #87 — $R^2$ = 0.15
Patient #88 — $R^2$ = 0.864
Patient #89 — $R^2$ = 0.931
Patient #90 — $R^2$ = 0.986
Patient #91 — $R^2$ = 0.965
Patient #92 — $R^2$ = 0.516
Patient #93 — $R^2$ = 0.46
Patient #94 — $R^2$ = -0.35
Patient #95 — $R^2$ = 0.691
Patient #96 — $R^2$ = 0.973

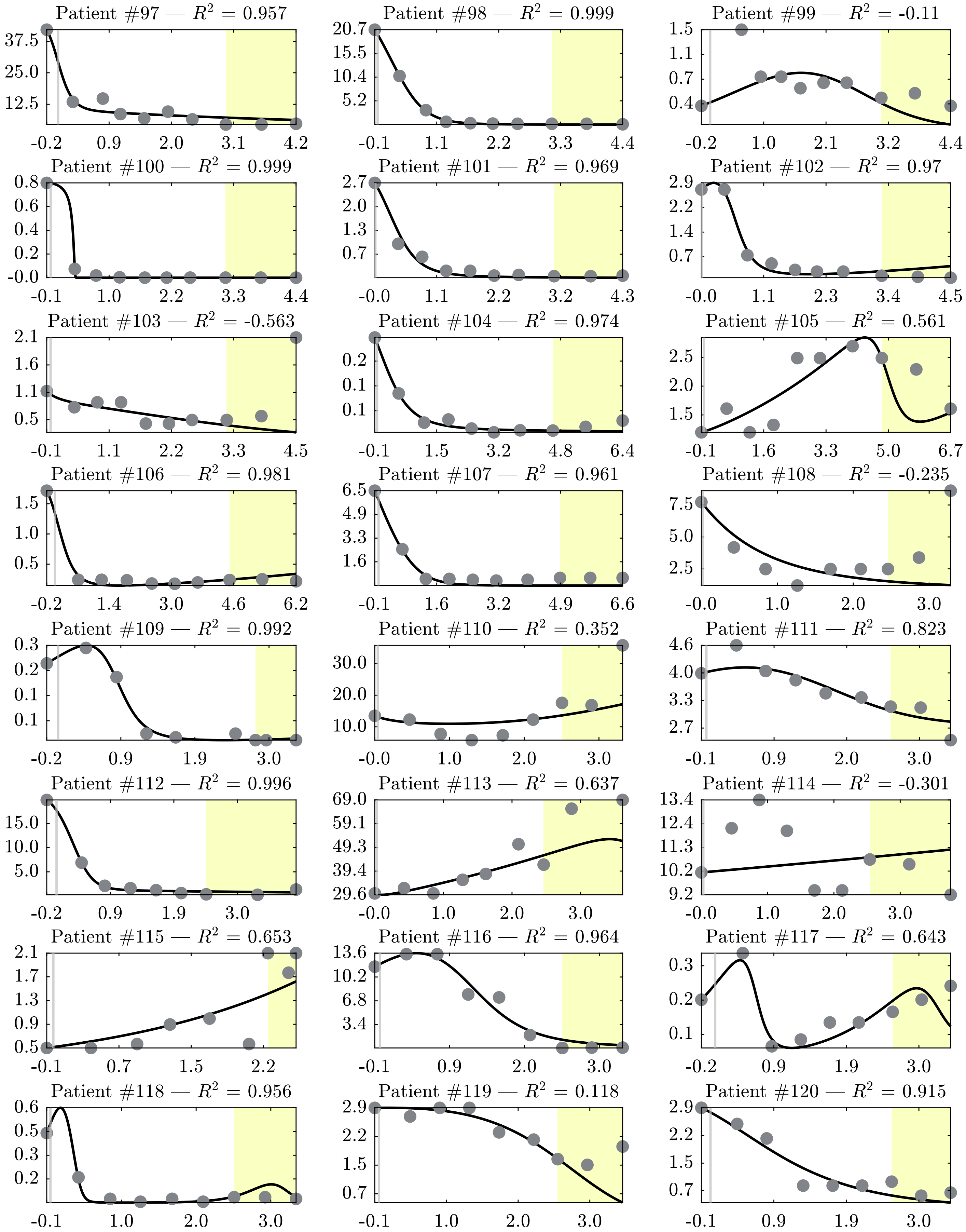

Patient #97 — $R^2$ = 0.957
37.5 25.0 12.5
-0.2 0.9 2.0 3.1 4.2
Patient #98 — $R^2$ = 0.999
20.7 15.5 10.4 5.2
-0.1 1.1 2.2 3.3 4.4
Patient #99 — $R^2$ = -0.11
1.5 1.1 0.7 0.4
-0.2 1.0 2.1 3.2 4.4
Patient #100 — $R^2$ = 0.999
0.8 0.6 0.4 0.2 -0.0
-0.1 1.0 2.2 3.3 4.4
Patient #101 — $R^2$ = 0.969
2.7 2.0 1.3 0.7
-0.0 1.1 2.2 3.2 4.3
Patient #102 — $R^2$ = 0.97
2.9 2.2 1.4 0.7
-0.0 1.1 2.3 3.4 4.5
Patient #103 — $R^2$ = -0.563
2.1 1.6 1.1 0.5
-0.1 1.1 2.2 3.3 4.5
Patient #104 — $R^2$ = 0.974
0.2 0.1 0.1
-0.1 1.5 3.2 4.8 6.4
Patient #105 — $R^2$ = 0.561
2.5 2.0 1.5
-0.1 1.6 3.3 5.0 6.7
Patient #106 — $R^2$ = 0.981
1.5 1.0 0.5
-0.2 1.4 3.0 4.6 6.2
Patient #107 — $R^2$ = 0.961
6.5 4.9 3.3 1.6
-0.1 1.6 3.2 4.9 6.6
Patient #108 — $R^2$ = -0.235
7.5 5.0 2.5
-0.0 1.0 2.0 3.0
Patient #109 — $R^2$ = 0.992
0.3 0.2 0.1 0.1
-0.2 0.9 1.9 3.0
Patient #110 — $R^2$ = 0.352
30.0 20.0 10.0
-0.0 1.0 2.0 3.0
Patient #111 — $R^2$ = 0.823
4.6 4.0 3.3 2.7
-0.1 0.9 2.0 3.0
Patient #112 — $R^2$ = 0.996
15.0 10.0 5.0
-0.2 0.9 1.9 3.0
Patient #113 — $R^2$ = 0.637
69.0 59.1 49.3 39.4 29.6
-0.0 1.0 2.0 3.0
Patient #114 — $R^2$ = -0.301
13.4 12.4 11.3 10.2 9.2
-0.0 1.0 2.0 3.0
Patient #115 — $R^2$ = 0.653
2.1 1.7 1.3 0.9 0.5
-0.1 0.7 1.5 2.2
Patient #116 — $R^2$ = 0.964
13.6 10.2 6.8 3.4
-0.1 0.9 2.0 3.0
Patient #117 — $R^2$ = 0.643
0.3 0.2 0.1
-0.2 0.9 1.9 3.0
Patient #118 — $R^2$ = 0.956
0.6 0.5 0.3 0.2
-0.1 1.0 2.0 3.0
Patient #119 — $R^2$ = 0.118
2.9 2.2 1.5 0.7
-0.1 1.0 2.0 3.0
Patient #120 — $R^2$ = 0.915
2.9 2.2 1.5 0.7
-0.1 0.9 1.9 3.0

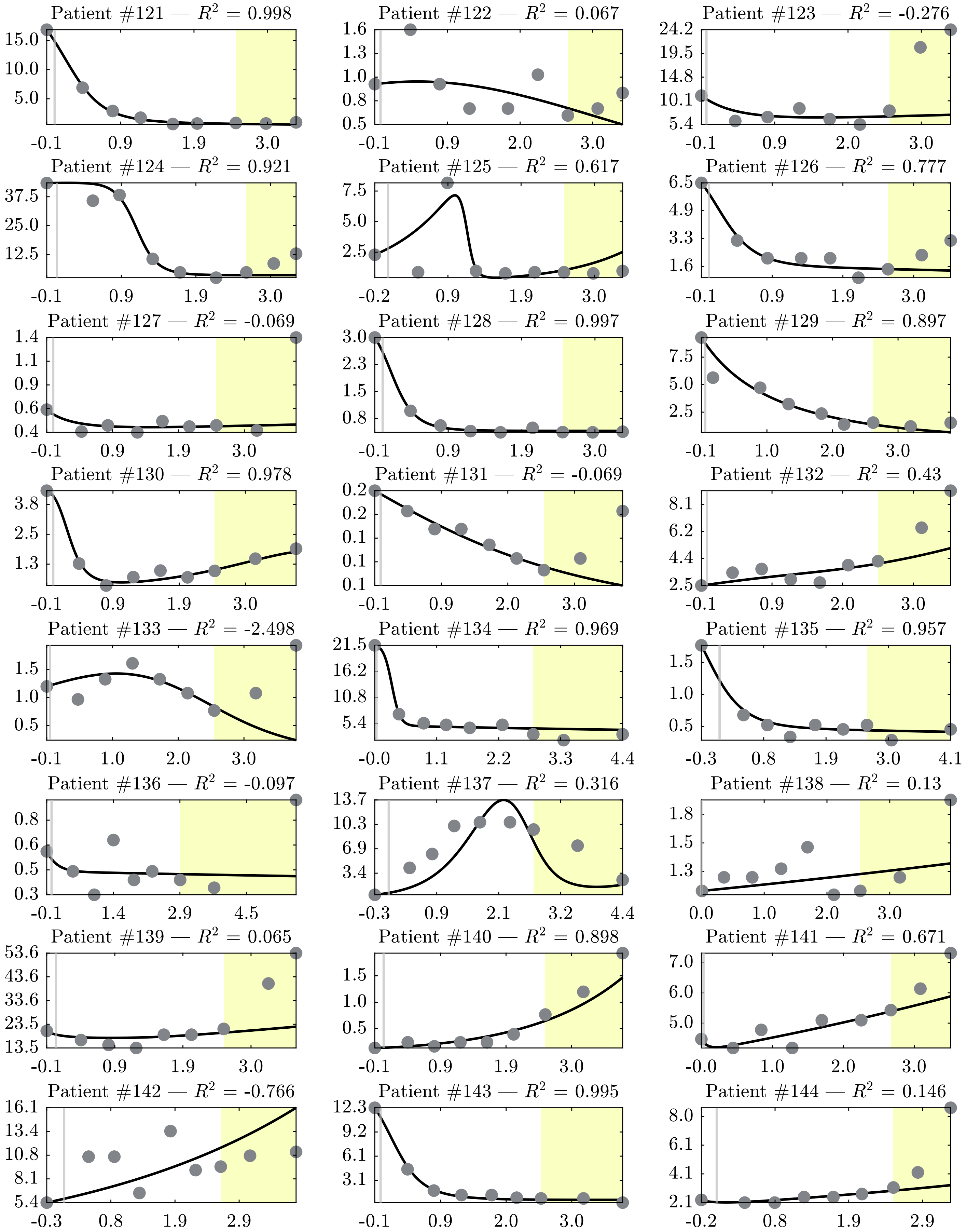
Patient #121 — $R^2$ = 0.998
15.0
10.0
5.0
-0.1
0.9
1.9
3.0
Patient #122 — $R^2$ = 0.067
1.6
1.3
1.0
0.8
0.5
-0.1
0.9
2.0
3.0
Patient #123 — $R^2$ = -0.276
24.2
19.5
14.8
10.1
5.4
-0.1
0.9
2.0
3.0
Patient #124 — $R^2$ = 0.921
37.5
25.0
12.5
-0.1
0.9
1.9
3.0
Patient #125 — $R^2$ = 0.617
7.5
5.0
2.5
-0.2
0.9
1.9
3.0
Patient #126 — $R^2$ = 0.777
6.5
4.9
3.3
1.6
-0.1
0.9
1.9
3.0
Patient #127 — $R^2$ = -0.069
1.4
1.1
0.9
0.6
0.4
-0.1
0.9
1.9
3.0
Patient #128 — $R^2$ = 0.997
3.0
2.3
1.5
0.8
-0.1
0.9
1.9
3.0
Patient #129 — $R^2$ = 0.897
7.5
5.0
2.5
-0.1
1.0
2.0
3.0
Patient #130 — $R^2$ = 0.978
3.8
2.5
1.3
-0.1
0.9
1.9
3.0
Patient #131 — $R^2$ = -0.069
0.2
0.2
0.1
0.1
0.1
-0.1
0.9
2.0
3.0
Patient #132 — $R^2$ = 0.43
8.1
6.2
4.4
2.5
-0.1
0.9
2.0
3.0
Patient #133 — $R^2$ = -2.498
1.5
1.0
0.5
-0.1
1.0
2.0
3.0
Patient #134 — $R^2$ = 0.969
21.5
16.2
10.8
5.4
-0.0
1.1
2.2
3.3
4.4
Patient #135 — $R^2$ = 0.957
1.5
1.0
0.5
-0.3
0.8
1.9
3.0
4.1
Patient #136 — $R^2$ = -0.097
0.8
0.6
0.5
0.3
-0.1
1.4
2.9
4.5
Patient #137 — $R^2$ = 0.316
13.7
10.3
6.9
3.4
-0.3
0.9
2.1
3.2
4.4
Patient #138 — $R^2$ = 0.13
1.8
1.5
1.3
0.0
1.0
2.0
3.0
Patient #139 — $R^2$ = 0.065
53.6
43.6
33.6
23.5
13.5
-0.1
0.9
1.9
3.0
Patient #140 — $R^2$ = 0.898
1.5
1.0
0.5
-0.1
0.9
1.9
3.0
Patient #141 — $R^2$ = 0.671
7.0
6.0
5.0
-0.0
1.0
2.0
3.0
Patient #142 — $R^2$ = -0.766
16.1
13.4
10.8
8.1
5.4
-0.3
0.8
1.9
2.9
Patient #143 — $R^2$ = 0.995
12.3
9.2
6.1
3.1
-0.1
0.9
2.0
3.0
Patient #144 — $R^2$ = 0.146
8.0
6.1
4.1
2.1
-0.2
0.8
1.9
2.9

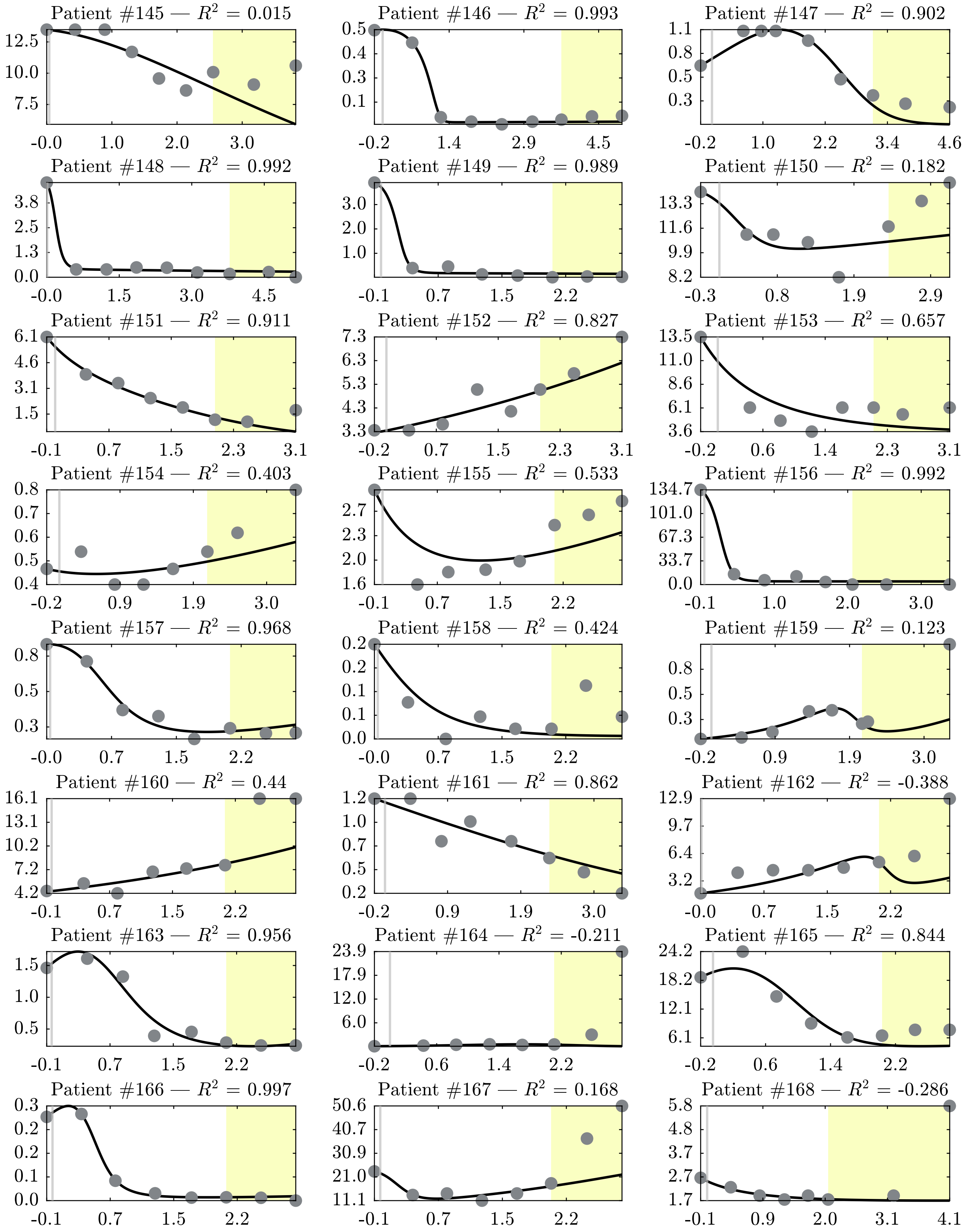

Patient #145 — $R^2$ = 0.015
12.5 10.0 7.5
-0.0 1.0 2.0 3.0
Patient #146 — $R^2$ = 0.993
0.5 0.4 0.3 0.1
-0.2 1.4 2.9 4.5
Patient #147 — $R^2$ = 0.902
1.1 0.8 0.5 0.3
-0.2 1.0 2.2 3.4 4.6
Patient #148 — $R^2$ = 0.992
3.8 2.5 1.3 0.0
-0.0 1.5 3.0 4.5
Patient #149 — $R^2$ = 0.989
3.0 2.0 1.0
-0.1 0.7 1.5 2.2
Patient #150 — $R^2$ = 0.182
13.3 11.6 9.9 8.2
-0.3 0.8 1.9 2.9
Patient #151 — $R^2$ = 0.911
6.1 4.6 3.1 1.5
-0.1 0.7 1.5 2.3 3.1
Patient #152 — $R^2$ = 0.827
7.3 6.3 5.3 4.3 3.3
-0.2 0.7 1.5 2.3 3.1
Patient #153 — $R^2$ = 0.657
13.5 11.0 8.6 6.1 3.6
-0.2 0.6 1.4 2.3 3.1
Patient #154 — $R^2$ = 0.403
0.8 0.7 0.6 0.5 0.4
-0.2 0.9 1.9 3.0
Patient #155 — $R^2$ = 0.533
2.7 2.3 2.0 1.6
-0.1 0.7 1.5 2.2
Patient #156 — $R^2$ = 0.992
134.7 101.0 67.3 33.7 0.0
-0.1 1.0 2.0 3.0
Patient #157 — $R^2$ = 0.968
0.8 0.5 0.3
-0.0 0.7 1.5 2.2
Patient #158 — $R^2$ = 0.424
0.2 0.2 0.1 0.1 0.0
-0.0 0.7 1.5 2.2
Patient #159 — $R^2$ = 0.123
0.8 0.5 0.3
-0.2 0.9 1.9 3.0
Patient #160 — $R^2$ = 0.44
16.1 13.1 10.2 7.2 4.2
-0.1 0.7 1.5 2.2
Patient #161 — $R^2$ = 0.862
1.2 1.0 0.7 0.5 0.2
-0.2 0.9 1.9 3.0
Patient #162 — $R^2$ = -0.388
12.9 9.7 6.4 3.2
-0.0 0.7 1.5 2.2
Patient #163 — $R^2$ = 0.956
1.5 1.0 0.5
-0.1 0.7 1.5 2.2
Patient #164 — $R^2$ = -0.211
23.9 17.9 12.0 6.0
-0.2 0.6 1.4 2.2
Patient #165 — $R^2$ = 0.844
24.2 18.2 12.1 6.1
-0.2 0.6 1.4 2.2
Patient #166 — $R^2$ = 0.997
0.3 0.2 0.2 0.1 0.0
-0.1 0.7 1.5 2.2
Patient #167 — $R^2$ = 0.168
50.6 40.7 30.9 21.0 11.1
-0.1 0.7 1.5 2.2
Patient #168 — $R^2$ = -0.286
5.8 4.8 3.7 2.7 1.7
-0.1 0.9 2.0 3.1 4.1

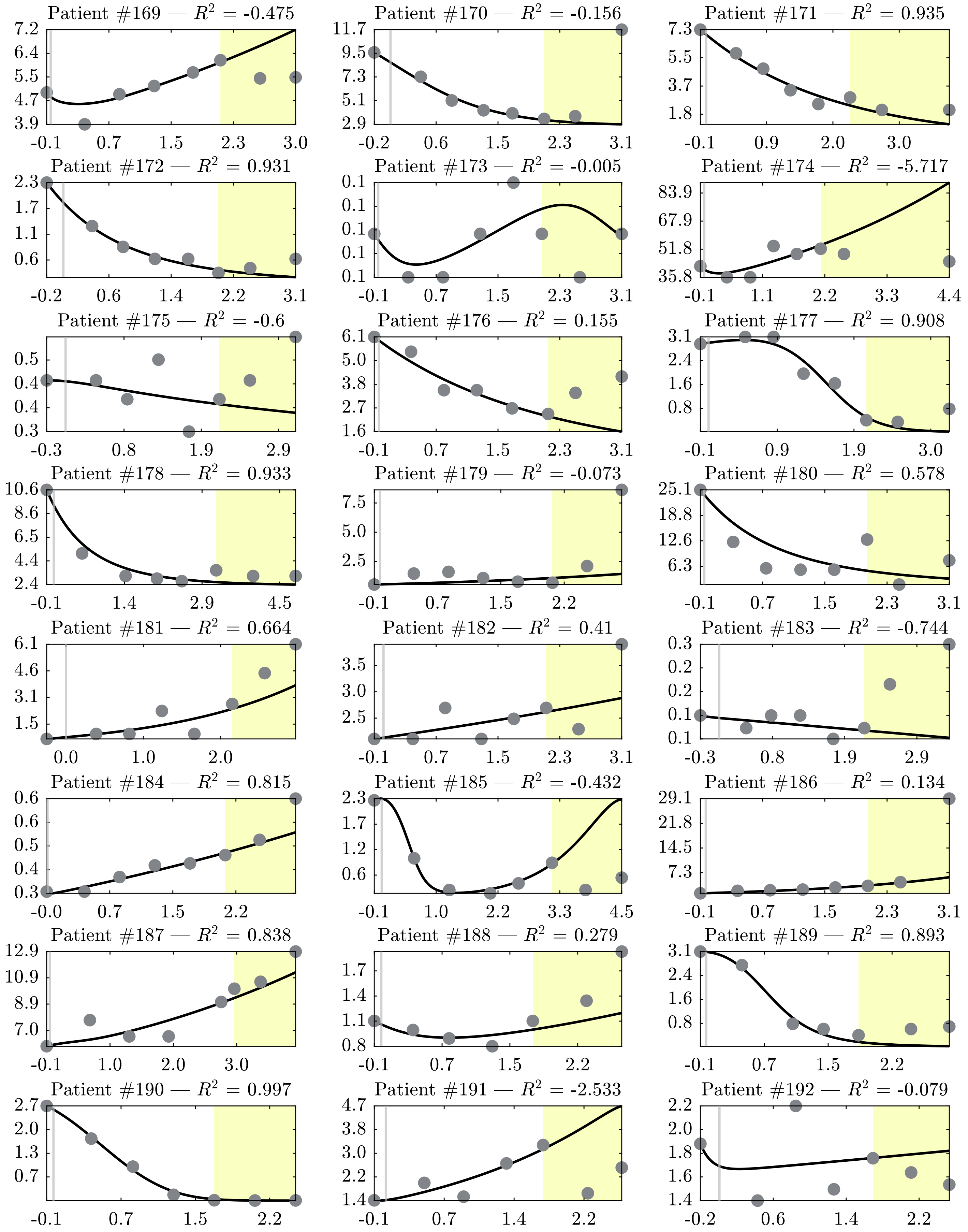
Patient #169 — $R^2$ = -0.475
7.2 6.4 5.5 4.7 3.9
-0.1 0.7 1.5 2.3 3.0
Patient #170 — $R^2$ = -0.156
11.7 9.5 7.3 5.1 2.9
-0.2 0.6 1.5 2.3 3.1
Patient #171 — $R^2$ = 0.935
7.3 5.5 3.7 1.8
-0.1 0.9 2.0 3.0
Patient #172 — $R^2$ = 0.931
2.3 1.7 1.1 0.6
-0.2 0.6 1.4 2.2 3.1
Patient #173 — $R^2$ = -0.005
0.1 0.1 0.1 0.1 0.1
-0.1 0.7 1.5 2.3 3.1
Patient #174 — $R^2$ = -5.717
83.9 67.9 51.8 35.8
-0.1 1.1 2.2 3.3 4.4
Patient #175 — $R^2$ = -0.6
0.5 0.4 0.4 0.3
-0.3 0.8 1.9 2.9
Patient #176 — $R^2$ = 0.155
6.1 5.0 3.8 2.7 1.6
-0.1 0.7 1.5 2.3 3.1
Patient #177 — $R^2$ = 0.908
3.1 2.4 1.6 0.8
-0.1 0.9 1.9 3.0
Patient #178 — $R^2$ = 0.933
10.6 8.6 6.5 4.4 2.4
-0.1 1.4 2.9 4.5
Patient #179 — $R^2$ = -0.073
7.5 5.0 2.5
-0.1 0.7 1.5 2.2
Patient #180 — $R^2$ = 0.578
25.1 18.8 12.6 6.3
-0.1 0.7 1.5 2.3 3.1
Patient #181 — $R^2$ = 0.664
6.1 4.6 3.1 1.5
0.0 1.0 2.0
Patient #182 — $R^2$ = 0.41
3.5 3.0 2.5
-0.1 0.7 1.5 2.3 3.1
Patient #183 — $R^2$ = -0.744
0.3 0.2 0.2 0.1 0.1
-0.3 0.8 1.9 2.9
Patient #184 — $R^2$ = 0.815
0.6 0.6 0.5 0.4 0.3
-0.0 0.7 1.5 2.2
Patient #185 — $R^2$ = -0.432
2.3 1.7 1.2 0.6
-0.1 1.0 2.2 3.3 4.5
Patient #186 — $R^2$ = 0.134
29.1 21.8 14.5 7.3
-0.1 0.7 1.5 2.3 3.1
Patient #187 — $R^2$ = 0.838
12.9 10.9 8.9 7.0
-0.1 1.0 2.0 3.0
Patient #188 — $R^2$ = 0.279
1.7 1.4 1.1 0.8
-0.1 0.7 1.5 2.2
Patient #189 — $R^2$ = 0.893
3.1 2.4 1.6 0.8
-0.1 0.7 1.5 2.2
Patient #190 — $R^2$ = 0.997
2.7 2.0 1.3 0.7
-0.1 0.7 1.5 2.2
Patient #191 — $R^2$ = -2.533
4.7 3.8 3.0 2.2 1.4
-0.1 0.7 1.4 2.2
Patient #192 — $R^2$ = -0.079
2.2 2.0 1.8 1.6 1.4
-0.2 0.6 1.4 2.2

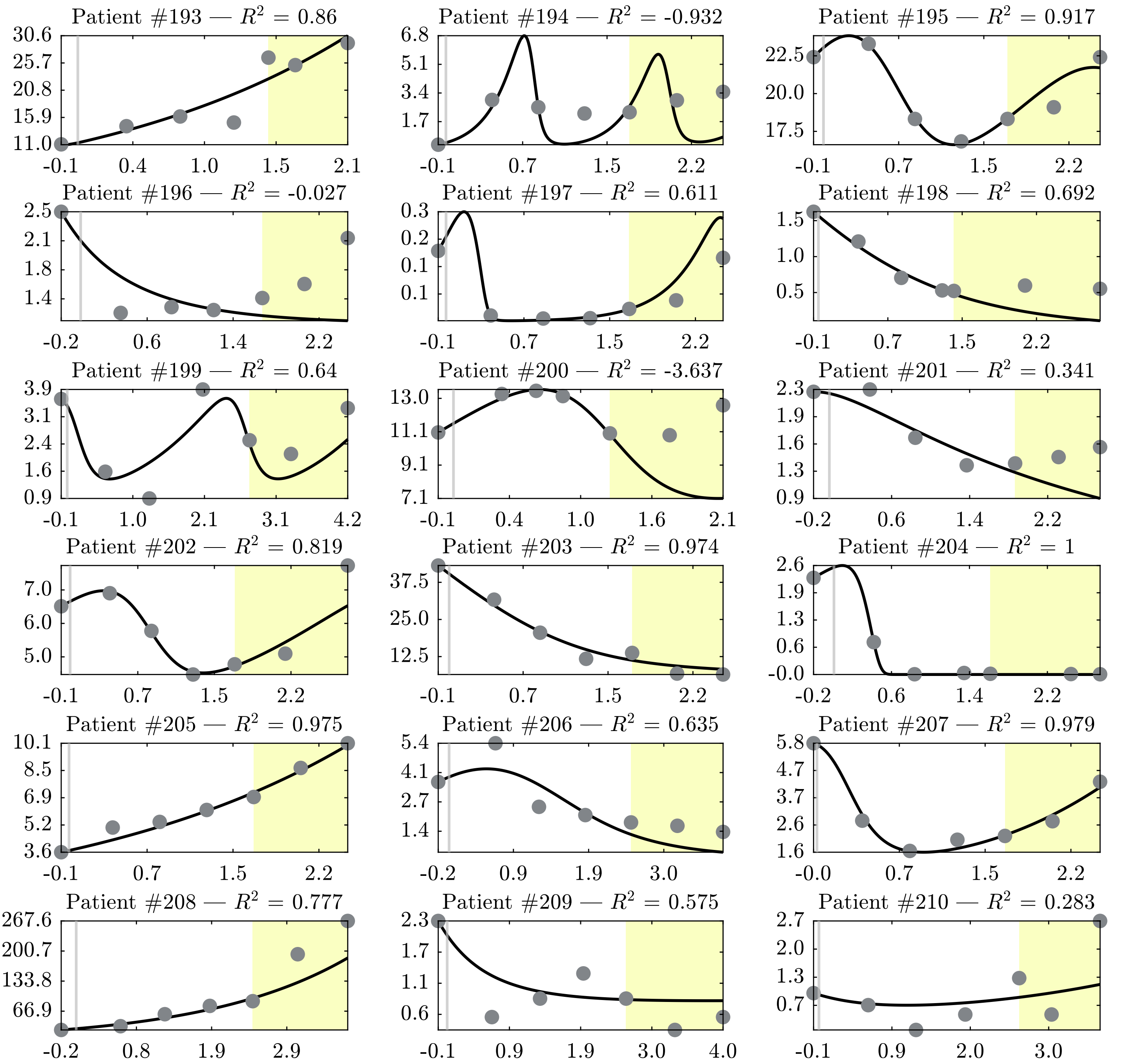
Patient #193 — $R^2$ = 0.86
30.6 25.7 20.8 15.9 11.0
-0.1 0.4 1.0 1.5 2.1
Patient #194 — $R^2$ = -0.932
6.8 5.1 3.4 1.7
-0.1 0.7 1.5 2.2
Patient #195 — $R^2$ = 0.917
22.5 20.0 17.5
-0.1 0.7 1.5 2.2
Patient #196 — $R^2$ = -0.027
2.5 2.1 1.8 1.4
-0.2 0.6 1.4 2.2
Patient #197 — $R^2$ = 0.611
0.3 0.2 0.1 0.1
-0.1 0.7 1.5 2.2
Patient #198 — $R^2$ = 0.692
1.5 1.0 0.5
-0.1 0.7 1.5 2.2
Patient #199 — $R^2$ = 0.64
3.9 3.1 2.4 1.6 0.9
-0.1 1.0 2.1 3.1 4.2
Patient #200 — $R^2$ = -3.637
13.0 11.1 9.1 7.1
-0.1 0.4 1.0 1.6 2.1
Patient #201 — $R^2$ = 0.341
2.3 1.9 1.6 1.3 0.9
-0.2 0.6 1.4 2.2
Patient #202 — $R^2$ = 0.819
7.0 6.0 5.0
-0.1 0.7 1.5 2.2
Patient #203 — $R^2$ = 0.974
37.5 25.0 12.5
-0.1 0.7 1.5 2.2
Patient #204 — $R^2$ = 1
2.6 1.9 1.3 0.6 -0.0
-0.2 0.6 1.4 2.2
Patient #205 — $R^2$ = 0.975
10.1 8.5 6.9 5.2 3.6
-0.1 0.7 1.5 2.2
Patient #206 — $R^2$ = 0.635
5.4 4.1 2.7 1.4
-0.2 0.9 1.9 3.0
Patient #207 — $R^2$ = 0.979
5.8 4.7 3.7 2.6 1.6
-0.0 0.7 1.5 2.2
Patient #208 — $R^2$ = 0.777
267.6 200.7 133.8 66.9
-0.2 0.8 1.9 2.9
Patient #209 — $R^2$ = 0.575
2.3 1.7 1.1 0.6
-0.1 0.9 1.9 3.0 4.0
Patient #210 — $R^2$ = 0.283
2.7 2.0 1.3 0.7
-0.1 0.9 2.0 3.0
